# CDK7, CDK9, or CDK11 Inhibition Reduces Neutrophil-Driven Inflammation and Tissue Damage in Experimental Autoimmune Models

**DOI:** 10.1101/2025.11.19.689221

**Authors:** Mareile Schlotfeldt, Leonie Voss, Adrian P. Mansini, Daniel Mehlberg, Sophia Johannisson, Michael Radziewitz, Ann-Kathrin Schneider, Seyed Mohammad Vahabi, Yulu Wang, Haley Gainer, Jing Li, Colin Osterloh, Nancy Ernst, Remco Visser, Gestur Vidarsson, Frank Petersen, Xinhua Yu, Ralf J. Ludwig, Artem Vorobyev, Kyle T. Amber, Anja Lux, Katja Bieber

## Abstract

**Background:** Cyclin-dependent kinases (CDKs) are involved in basic cellular processes like regulation of cell-cycle progression and transcription. However, recent data also indicate a specific role in terminally differentiated neutrophils by promoting reactive oxygen species (ROS) release, degranulation, neutrophil extracellular trap formation, or apoptosis. Since these mechanisms are implicated in multiple autoimmune diseases, we aimed to delineate the role of CDKs in IC-mediated autoimmune diseases both *in vitro* and *in vivo*.

**Methods:** We analyzed CDK gene expression in unstimulated and immune complex (IC)-stimulated neutrophils. Subsequently, we investigated the effect of pharmacological CDK inhibition on IC-activated neutrophil functions. To analyze the inhibitors in a more translational approach, we proceeded with the systemic and topical application of the effective inhibitors in a murine antibody transfer-induced local epidermolysis bullosa acquisita (EBA) model. The most efficient inhibitor, MC180295, was validated in two other IC-mediated models of autoimmune disease: Serum-transfer arthritis (STA), which also, but not exclusively, depends on neutrophils and immune thrombocytopenia (ITP), which is considered less neutrophil-dependent.

**Results:** We found 14 CDKs expressed in unstimulated cells, while the IC-stimulation showed an upregulation of CDK2 and CDK4 expression. Inhibitors selectively targeting CDK1, CDK2, CDK4/6, CDK7, CDK9, CDK11, and CDK12 showed effects on different neutrophil functions (surface activation marker expression, ROS release, adhesion, apoptosis) *in vitro*. In the predominantly neutrophil-driven EBA model, we observed a reduction of disease severity upon treatment with CDK7, CDK9, or CDK11 inhibitors. Inhibiting these CDKs with topical THZ2, MC180295, or OTS964, respectively, also improved the clinical phenotype. In line with our hypothesis, MC180295 impaired the development of STA, but not ITP.

**Conclusions:** CDK7, CDK11, and especially CDK9 inhibition **s**how therapeutic potential in IC-driven neutrophil-mediated diseases such as rheumatoid arthritis and EBA.

## Background

Cyclin-dependent kinases (CDKs) are a group of serine-threonine kinases that were discovered for their involvement in the regulation of the eukaryotic cell cycle [1–3]. CDKs are crucially contributing to the correct replication and subsequent separation of the chromosomal DNA during mitosis [4] and are highly conserved across eukaryotes, from yeast to mammals [5]. They were termed for their critical interaction with specific regulatory protein binding partners, the cyclins [6]. The latter are expressed in a cyclic fashion during the cell cycle, while the CDKs themselves are stably expressed throughout the cell cycle [3]. There are 20 CDKs and 29 cyclins described in mammals [7], as well as other regulating proteins such as intrinsic CDK inhibitors (CKI) [8]. Mechanistically, activation of CDKs not only requires the interaction with a cyclin, also phosphorylation at a specific tyrosine residue in the activation loop is necessary [9, 10]. This phosphorylation of at least CDK1/2/4/6/9/12/13 is mediated by the CDK-activating kinase (CAK) complex [11], consisting of CDK7, cyclin H and MAT1 [12–14]. Importantly, CDK inactivation is also regulated by phosphorylation [15] or binding of CKIs [16, 17].

Beyond the well-described role of CDKs in direct cell-cycle regulation, there is rising evidence that CDKs are also involved in regulating the RNA polymerase II transcription cycle [11, 18]. It was thus proposed that there are two groups of CDKs: the cell-cycle CDKs (subfamilies CDK1, 2, 3, 4, and 6) and the transcriptional CDKs (subfamilies CDK7, 8, 9, 10, 11, 12, and 13) [7]. The remainder is referred to as atypical CDKs, although these kinases are insufficiently characterized [19]. Though this may be a helpful concept, this dichotomy is probably oversimplified as more and more studies demonstrate roles for traditional cell-cycle CDKs in transcriptional regulation and vice versa [18, 20–22].

Considering that CDKs are not only expressed in proliferating cells, but also in terminally differentiated cells like neutrophils [23], it seems a reasonable assumption that their role extends beyond mere cell cycle regulation. Neutrophils are known as the first line of defense against pathogens [24]. Once recruited to the tissue of interest, they are activated by the binding of immune complexes (IC) to their fragment crystallizable gamma receptors (FcγR). Downstream kinase signaling cascades are activated, triggering effector functions such as reactive oxygen species (ROS), protease and cytokine release to kill the invading pathogens in a controlled manner. Subsequently, the neutrophils are cleared mostly by macrophages under inflammatory conditions [25, 26]. Notably, a role for several CDKs in regulating neutrophil functions has been described: CDK2 is involved in migration [27], CDK4/6 in formation of neutrophil extracellular traps (NETs) [28], CDK5 in degranulation [29], and CDK7/9 in apoptosis [22, 30]. Still, little is known about the distinct functions of different CDKs in direct comparison, and most studies rely on broad-range small-molecule CDK inhibitors (CDKi). Moreover, the potential involvement of CDKs in FcγR-mediated signaling remains largely unexplored, representing a significant gap in our understanding of CDK biology.

An increasing number of studies points towards the relevance of CDKs in various human disorders, underlining the need for a better comprehension of these kinases. Deregulated CDK activity has been shown to be associated not only with e.g. neurodegenerative diseases [31] and cancer [19], but also with autoimmune diseases like rheumatoid arthritis (RA) [32] or systemic lupus erythematosus [33]. In contrast to infectious diseases, however, there is no pathogen in autoimmunity, but autoantibodies develop against self-antigens due to a break of tolerance [34]. To unravel the function of CDKs in the context of neutrophil-mediated FcγR-dependent signaling, we investigated the kinase signaling pathways upon IC stimulation in human neutrophils *in vitro.* Furthermore, we chose three well-established murine *in vivo* models of antibody-mediated autoimmune diseases: Epidermolysis bullosa acquisita (EBA), serum-transfer arthritis (STA) as a model for RA, and immune thrombocytopenia (ITP). EBA belongs to the group of pemphigoid diseases (PD), which is a group of rare autoimmune bullous skin diseases characterized by autoantibodies targeting structural proteins of the dermal-epidermal junction. More specifically, the autoantigen in EBA is collagen VII (COL7) [35]. Neutrophils are the central effector cells in PD [36–39], causing the uncontrolled release of inflammatory mediators. Thereby, tissue damage occurs, causing the degradation of the dermal-epidermal adhesion complex and subepidermal split formation [38, 40]. Clinically, patients present with pruritus, erosions, and tense, painful blisters on different sites of the body, e.g. trauma-prone areas or the extensor skin surface [41]. Currently, the patients are most commonly treated with systemic corticosteroids, often in combination with additional immunosuppressants, bearing a high risk for adverse events due to general immunosuppression [42, 43].

Another more prevalent neutrophil-dependent autoimmune disease with the need for novel therapeutic options is RA, modelled by STA [44]. In contrast to EBA, neutrophils do play a critical role in pathogenesis [45, 46], but other cell types like macrophages [47] or mast cells [48, 49] are crucially involved in the effector phase. Joint inflammation, bone and cartilage destruction are major hallmarks of RA [50].

Conversely, ITP is also IC-driven, but depends less on neutrophils. In ITP, autoantibodies target platelets leading to their phagocytosis mainly by macrophages and thus to platelet depletion [51–53].

Since kinases are generally well-druggable targets [54, 55] and considering that there are several CDKi already in clinical use, e.g. for cancer treatment [56], a potential repurposing of the inhibitors could accelerate the process of clinical approval. Thus, the characterization of CDK signaling cascades in neutrophils can help to identify potential therapeutic targets for PD, but also potentially for other neutrophil-driven autoimmune diseases like RA. This could provide targeted treatment options to the patients and therefore drastically ameliorate the safety profile of the therapy.

## Methods

### Sampling of human biomaterials

Whole blood was obtained from healthy volunteers after written informed consent. All experiments using human samples were approved by the local ethics committee (AZ #20-463, University of Lübeck, Lübeck, Germany) and were performed in accordance with the Declaration of Helsinki.

### Chemicals

All standard chemicals were supplied by Carl Roth (Karlsruhe, Germany) or Sigma-Aldrich (Taufkirchen, Germany). CDKi were supplied by Selleck Chemicals (Houston, Texas, USA) or MedChemExpress (Monmouth Junction, New Jersey, USA). For *in vitro* assays, 10 mM CDKi was dissolved in dimethyl sulfoxide (DMSO) and further diluted to final concentrations of 10 µM, 1 µM, 0.1 µM, and 0.01 µM. DMSO at a final concentration of 0.1 % was included in the controls. For *in vivo* experiments, CDKi were prepared as stated in Table 1. The respective vehicle only served as the respective control. Additional targets, references and stage of drug development are given in Supplemental table 1.

**Table 1.**
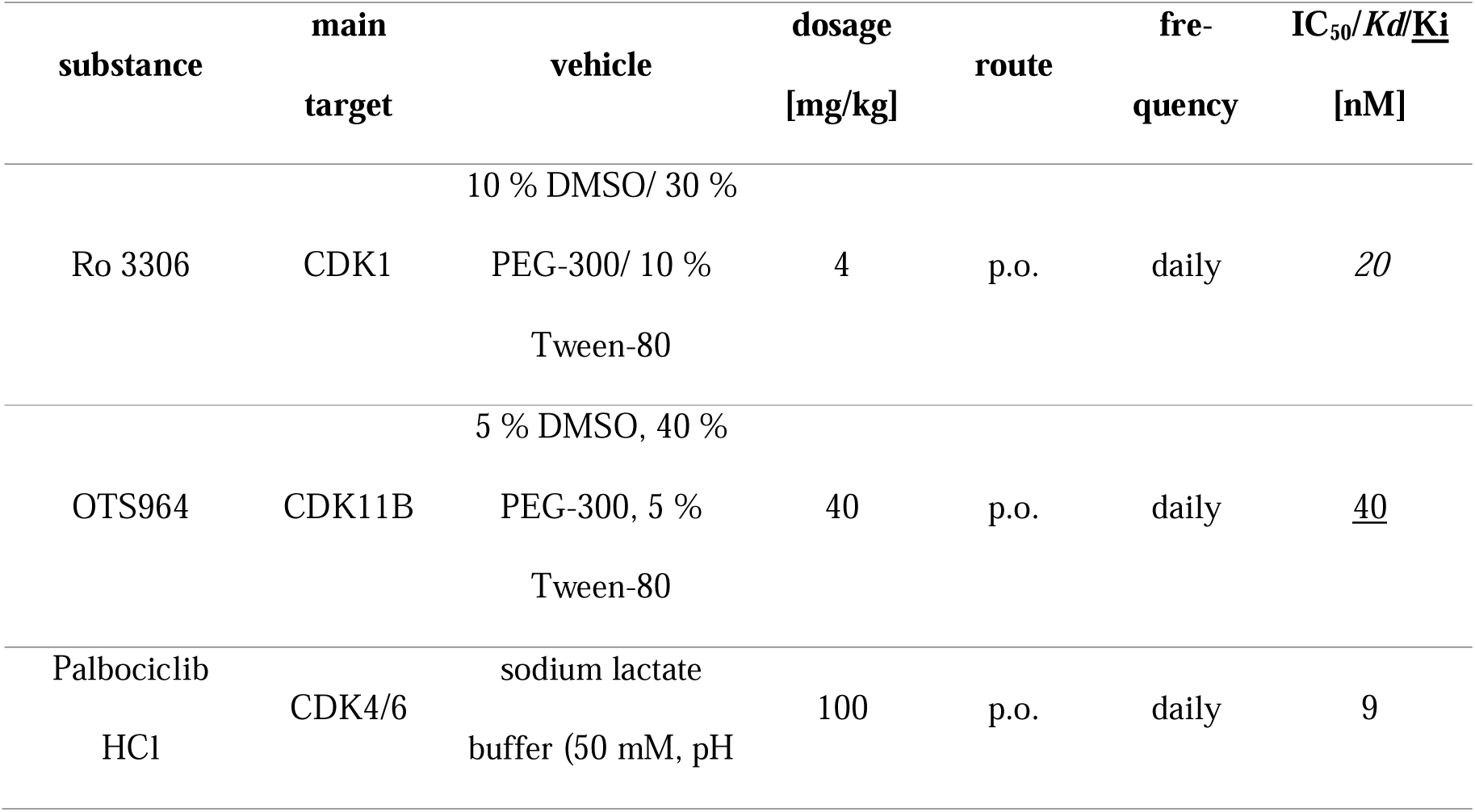
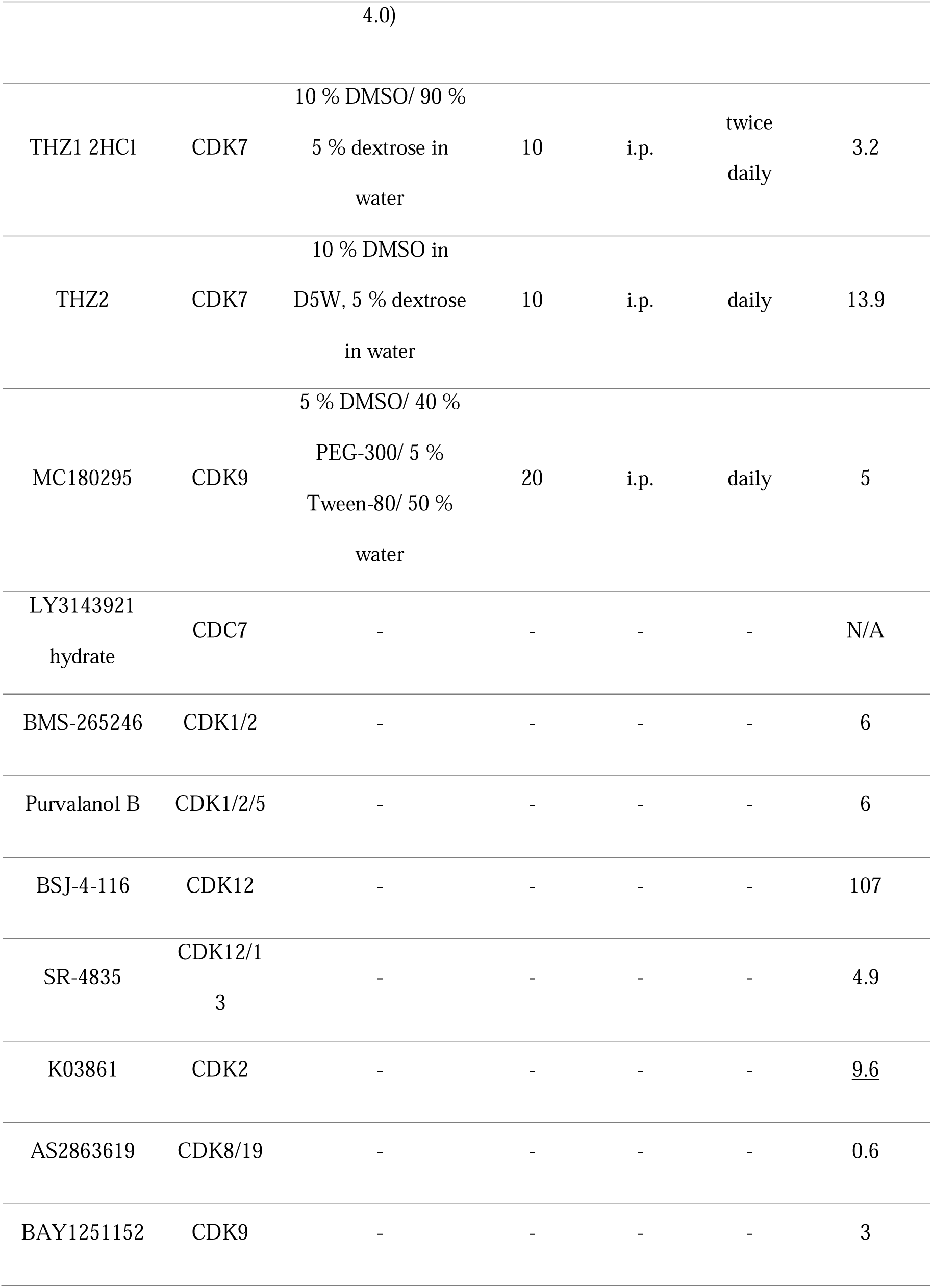
CDKi used in this study. The vehicle, dosage, route and frequency for *in vivo* treatment stated, where applicable. IC_50_, Kd, or Ki values were adapted from Selleckchem.com. N/A, not available.

**Supplemental table 1.**
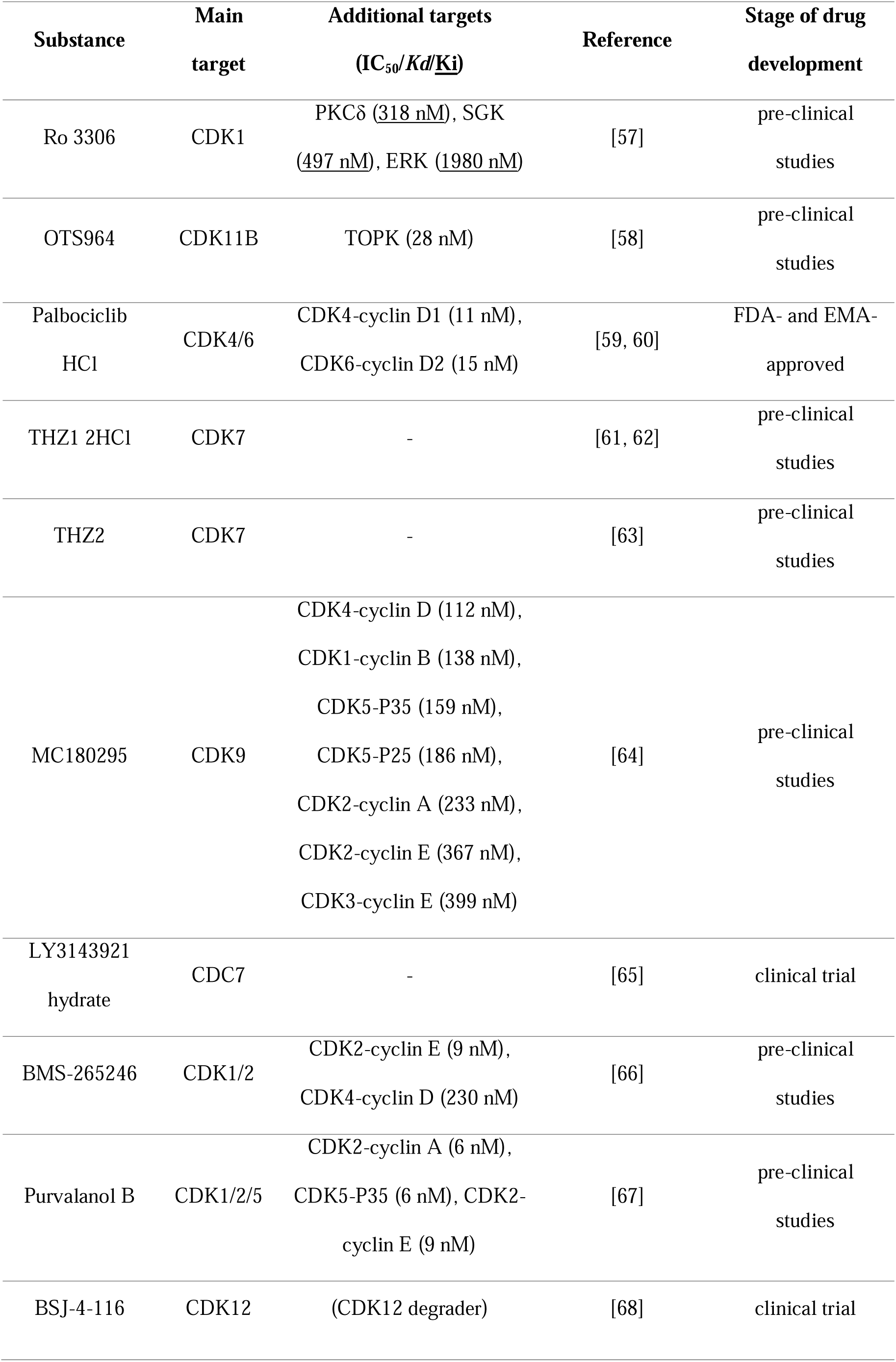
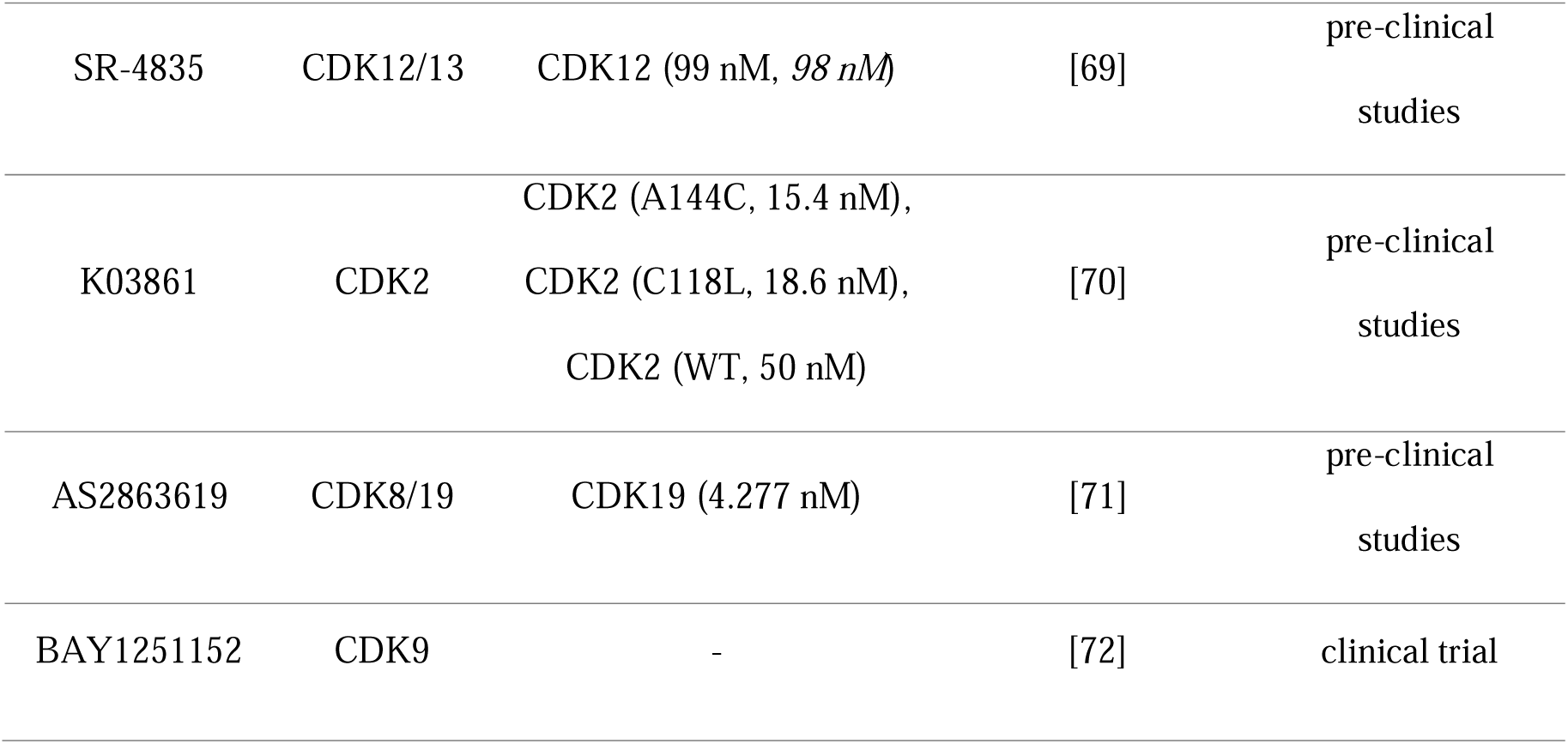
Additional targets of CDKi used adopted from Selleckchem.com and references for *in vivo* application and current stage of drug development.

### Isolation of human immune cells by magnetic activated cell sorting (MACS)

For the isolation of human neutrophils by MACS, blood from healthy young (< 30 y) donors was drawn into citrate blood collection tubes. After 1:2 dilution with 1 % (w/v) polyvinyl alcohol and sedimentation for 25 min, the supernatant (up to 40 mL) was transferred on top of a layer of 8 mL of Ficoll Paque PLUS (Cytiva, Marlborough, Massachusetts, USA). Density gradient centrifugation [73] was performed for 24 min at 850 g and room temperature (RT) without brake. The lymphocyte layer was processed for B and CD4^+^ T cell isolation, whereas the granulocyte pellet was used for neutrophil purification. The lymphocyte layer was resuspended in phosphate-buffered saline (PBS) and centrifuged at 300 g for 10 min at 4 °C for platelet removal. Another washing step with PBS was performed at 400 g for 10 min at 4 °C. Next, the purified lymphocytes were subjected to MACS according to the manufacturer’s instructions using the CD4^+^ T cell isolation kit (130-096-533, Miltenyi Biotec, Bergisch Gladbach, Germany) and the B cell isolation kit II (130-091-151, Miltenyi Biotec). B and CD4^+^ T cell purities were assessed by flow cytometry and were > 80 %.

For erythrocyte lysis, the granulocyte pellet was resuspended in distilled water, mixed for 45 s, and then neutralized with 2x PBS. Two washing steps with PBS were performed for 10 min at 300 g and RT. Subsequently, the granulocytes were subjected to eosinophil removal by MACS according to the manufacturer’s instructions (130-092-010, Miltenyi Biotec). Neutrophil purity was assessed by flow cytometry and was > 98 %.

### Bulk RNA sequencing

For bulk RNA sequencing (RNAseq), 100,000 MACS-purified neutrophils from healthy volunteers were sampled and frozen at -80 °C in 1 mL Trizol (Invitrogen, Waltham, Massachusetts, USA). An equal number of cells from each donor was stimulated for 6 h at 37 °C and 5 % CO_2_ with immobilized IC consisting of human COL7E-F and anti-human COL7 IgG1 as previously described [74]. All samples were sent to Single Cell Discoveries (Utrecht, The Netherlands) for RNA extraction and sequencing. Data analysis was performed in RStudio (Posit Software, Boston, Massachusetts, USA; version 2024.12.1) as previously described [75]. Differentially expressed genes were identified using DESeq2 [76] on transcripts where the sum of the counts over all samples was > 10,and p_adj_ < 0.1 was considered significant. Gene set enrichment analysis was conducted using ClusterProfiler referring to the Gene Ontology (GO) database [77, 78]. The data for this study have been deposited in the European Nucleotide Archive (ENA) at EMBL-EBI under accession number PRJEB103987 (https://www.ebi.ac.uk/ena/browser/view/PRJEB103987).

### Human polymorphonuclear granulocyte (PMN) purification for real-time quantitative reverse transcription polymerase chain reaction (qRT-PCR) and functional assays

For ROS release assay, flow cytometry, adhesion assay, and qRT-PCR, PMNs were isolated from EDTA-treated human whole blood performing a PolymorphPrep™ (Serumwerk Bernburg AG, Bernburg, Germany) density gradient centrifugation as previously described [74, 75, 79]. The cells were diluted to 4×10^5^ cells/mL in 1 % (w/v) bovine serum albumin (BSA) in PBS. Neutrophil purity was assessed by flow cytometry for all experiments and was higher than 90 % for all samples.

### qRT-PCR

For gene expression analysis by qRT-PCR, human PMNs were stimulated with human COL7E-F anti-human COL7 IgG1 IC for 6 h at 37 °C and 5 % CO_2_ as stated above. Alongside unstimulated PMNs, B cells and CD4^+^ T cells isolated by MACS as described above, the cells were collected with a cell scraper in PBS and centrifuged for 7 min at 700 g and RT. The lysates were resuspended in 700 µL RLT buffer (Qiagen, Hilden, Germany) and frozen at -80 °C until RNA extraction. RNA extraction was performed using the innuPREP RNA Mini Kit 2.0 (Analytik Jena, Jena, Germany) according to the manufacturer’s instructions. Any remaining RNA was digested with DNase I (Sigma-Aldrich). cDNA was synthesized using the RevertAid Reverse Transcriptase (Thermo Fisher Scientific, Waltham, Massachusetts, USA) according to the manufacturer’s instructions. TaqMan™ gene expression assays (see Table 2) were purchased from Thermo Fisher Scientific and used according to the manufacturer’s instructions. qRT-PCR was performed on an Applied Biosystems 7900HT Fast Real-Time PCR System (Thermo Fisher Scientific). The results were analyzed using the 2^-ΔΔCt^ method [80] and normalized to the mean values of the housekeeping genes [81].

**Table 2.** TaqMan™ gene expression assays used for qRT-PCR.

| CDKs | Catalogue<br>no. | Assay ID | Gene symbol |
| --- | --- | --- | --- |
|  | 4453320 | Hs00938777_m1 | *CDK1* |
|  | 4453320 | Hs01548894_m1 | *CDK2* |
|  | 4448892 | Hs01077186_g1 | *CDK3, TEN1-CDK3* |
|  | 4453320 | Hs00364847_m1 | *CDK4* |
|  | 4453320 | Hs00358991_g1 | *CDK5* |
|  | 4453320 | Hs01026371_m1 | *CDK6* |
|  | 4448892 | Hs00361486_m1 | *CDK7* |
|  | 4448892 | Hs00992501_g1 | *CDK8* |
|  | 4448892 | Hs00977896_g1 | *CDK9* |
|  | 4448892 | Hs02341397_m1 | *CDK11A* |
|  | 4448892 | Hs07291776_gH | *CDK11B* |
|  | 4448892 | Hs00901732_m1 | *CDK12* |
|  | 4448892 | Hs00953418_m1 | *CDK14* |
|  | 4448892 | Hs01039930_g1 | *CDK19* |
|  | 4453320 | Hs00177487_m1 | *CDC7* |
| housekeeping genes | 4331182 | Hs01060665_g1 | *ACTB* |
|  | 4331182 | Hs02786624_g1 | *GAPDH* |
|  | 4331182 | Hs02800695_m1 | *HPRT* |
|  | 4331182 | Hs00272002_m1 | *GNB2L1* |

### ROS release assay

Plates were coated with IC consisting of human COL7 E-F at 2.5 µg/mL and anti-human COL7 IgG1 at 1.8 µg/mL as previously described [74, 82]. PMNs were added at 2×10^5^ cells per well in the presence or absence of CDKi. Only antibody (with cells), only antigen (with cells), or only cells served as negative controls, while IC (with cells, but without CDKi) served as positive control. DMSO was adjusted to a final concentration of 0.01 %. All samples were measured in duplicates. Luminol was automatically injected into each well to reach a final concentration of 90 µg/mL, and chemiluminescence was measured for 2 h every 2 min at 37 °C in a GloMax® Discover microplate reader (Promega, Walldorf, Germany). The area under the curve (AUC) was calculated, and the results were normalized to IC-stimulated cells without CDKi.

### Adhesion assay

PMN adhesion was investigated using impedance-based measurements as described before [83, 84]. A 96-well E-plate (Agilent, Santa Clara, California, USA) was coated with IC as described for the ROS release assay. Inhibitor solutions were prepared and added to the plate before 2×10^5^ cells were transferred to each well. Only antibody (with cells), only antigen (with cells), or only cells served as negative controls, while IC (with cells, but without CDKi) served as positive control. DMSO was adjusted to a final concentration of 0.01 %. All samples were measured in duplicates. Impedance was measured in an xCELLigence real-time cell analysis device (Agilent) every 2 min for 120 min expressed in arbitrary units as cell index. Data analysis was performed as for the ROS release assay.

### Flow cytometry

Human PMNs were isolated as previously described and stimulated with IC, prepared in the same manner as for the ROS release assay, for 2 h at 37_°C and 5 % CO_. Subsequently, they were stained for the flow cytometric analysis of the following surface markers using standard flow cytometry procedures: CD14 (clone HCD14), CD16 (clone 3G8), CD18 (clone 1B4/CD18), CD45 (clone HI30), CD62L (clone DREG-56), and CD193 (clone 5E8). Viability was assessed by Annexin V (B266195) and Zombie NIR staining (B308933) [75]. Only antibody (with cells), only antigen (with cells), or only cells served as negative controls, while IC (with cells, but without CDKi) served as positive control. DMSO was adjusted to a final concentration of 0.01 %. All samples were measured in duplicates. Analyses for activation markers were performed on FSC-SSC gated CD45^+^ singlets. Annexin V^-^ Zombie NIR^-^ cells were considered viable, Annexin V^+^ Zombie NIR^-^ cells were considered apoptotic. The positive control corresponds to values of up to 3 %. All dyes were purchased from BioLegend (San Diego, California, USA). Measurements were performed on a MACSQuant® Analyzer 10 (Miltenyi Biotec) and analysis was conducted using MACS Quantify software (version 2.13.3, Miltenyi Biotec).

### Study approval

Animal experiments were approved by local authorities of the Animal Care and Use Committee (government of Schleswig-Holstein and government of Lower Franconia) and the Institutional Animal Care and Use Committee at Rush University. All experiments were performed by certified personnel (AZ 108/08-15, AZ 97-11/20 and 55.2.2-2532-2-2085) following the ARRIVE guidelines.

### Animal experimentation

C57BL/6J mice (Charles River, Wilmington, Massachusetts, USA) were bred in a specific pathogen–free environment and provided standard mouse chow and acidified drinking water *ad libitum*. Mice of both sexes, aged 8 to 14 weeks, were used for experimental EBA models with clinical examinations performed under anaesthesia, using intraperitoneal (i.p.) administration of a ketamine (75 mg/kg, Sigma-Aldrich) and medetomidine (1 mg/kg, Vetoquinol, Ismaning, Germany) mixture, antagonized after ca. 30 min by atipamezole (5 mg/kg, Vetoquinol). On the final day, the mice were anaesthetized using 15 mg/kg xylazine (Sigma-Aldrich) and 100 mg/kg ketamine. STA and ITP models were performed with female C57BL/6J aged 12 weeks. For anaesthesia, isoflurane was used and mice were killed by CO_2_.

### Local antibody-transfer induced experimental EBA

Specific anti–mouse COL7^C^ IgG from rabbit serum was isolated as previously described [40, 85–87]. For systemic treatment, CDKi were administered as indicated in Table 1 at 100 µL/20 g mouse. For topical treatment in independent experiments, THZ2, MC180295, or OTS964 were applied at 65 µg/100 µL in acetone/DMSO (20:1) per ear, beginning one day before the anti–mouse COL7^C^ IgG injection and treatment was performed daily for four days. Vehicle-treated mice served as controls in all cases. Experimental EBA was induced by ear-base injection with 100 μg rabbit anti–mouse COL7^C^ IgG. The mice were clinically evaluated every day until day 3, and the percentage of ear thickness was assessed using a Mitutoyo 7301 dial thickness gauge (Neuss, Germany) by a blinded person, as well as the affected ear surface area (AESA). Ears were collected on the final day of the experiment, and half of each ear was embedded in 4 % PBS-buffered paraffin or frozen in liquid nitrogen and stored at -80 °C. Histological analysis of hematoxylin and eosin (H&E) staining and direct immunofluorescence staining for murine IgG and C3 were performed as described elsewhere [87]. Scoring of H&E-stained sections was performed to determine (i) the magnitude of epidermal thickness with values from 0 (10-20 µm), 1 (20-40 µm), 2 (40-100 µm) to 3 (> 100 µm), (ii) the percentage of dermal-epidermal split formation (length of split formation/length of dermal-epidermal junction), with values from 0 (no split), 1 (< 20 %), 2 (20-50 %) to 3 (> 50 %) and (iii) the dermal infiltration with values from 0 to 3 corresponding to no, mild, moderate or severe degree, respectively. The H&E score was calculated as the mean of these three parameters.

### STA

STA was induced by injection of arthritogenic KBxN serum in female C57BL/6J mice at 10 µL/g body weight as described before [88]. Before and after arthritis induction, mice were treated daily with 20 mg/kg body weight MC180295 or vehicle control i.p. The severity of joint swelling was monitored to evaluate clinical disease over the course of seven days. Peripheral blood was collected retro-orbitally on day 0 and day 7, and inflammatory cytokines were quantified in the serum by LegendPlex multiplex assay (Mouse Inflammation 13-plex, BioLegend) as a parameter of systemic inflammation. On day 7, mice were sacrificed, and forelegs were prepared for histological analysis. H&E staining was performed to evaluate inflammatory infiltrates and Safranin-O staining to assess bone and cartilage damage, as previously described [88, 89]. Stained sections were evaluated by light microscopy using a Zeiss Axiovert 200 microscope and the AxioVision LE4.6 software (Olympus Life Science, Waltham, Massachusetts, USA).

### Immune thrombocytopenia (ITP)

Platelet depletion in the presence of kinase inhibition was assessed in female C57BL/6J mice. Platelet depletion was induced by injection of 0.35 µg/g anti-platelet IgG2c (clone 6A6) as described [53]. Treatment with vehicle control or MC180295 (20 mg/kg body weight i.p.) was performed 24 h and 4 h before the induction of platelet depletion. Platelet counts in peripheral blood were assessed immediately before 6A6-IgG2c injection, and at 4 h and 24 h post-injection in peripheral blood collected retro-orbitally. Platelet counts were determined using an ADVIA® 2120i hematocytometer (Siemens Healthineers, Erlangen, Germany).

### Statistical analysis

Data were analyzed using Prism 10.4.1 (GraphPad Software, San Diego, USA). For all *in vitro* assays, Kruskal-Wallis test with Dunn’s multiple comparisons test was performed if not indicated differently. Dose response curves and IC_50_ values were calculated using the non-linear fit inhibitor vs. response (three parameters) function or for EC_50_ values using the agonist vs. response (three parameters) function where an activating response was observed. For all *in vivo* experiments, two-way ANOVA or mixed-effects analysis (in the case of missing values) with Šidák’s multiple comparisons test were performed unless stated differently. A value of p ≤ 0.05 was considered significant. The data are displayed as Tukey box-and-whisker plots or else as mean ± standard deviation unless described differently.

## Results

### Fourteen CDKs are expressed at mRNA level in human neutrophils and expression of CDK2 and CDK4 changes upon IC stimulation

CDKs are essential to drive cell cycle progression in proliferating cells [19]. However, at least certain CDKs seem to be functionally relevant also in terminally differentiated cells like neutrophils [22, 27–29]. Since there are conflicting data published on this topic [22, 28], we first set out to clarify which CDKs and regulating proteins are expressed at mRNA level in human neutrophils. Therefore, an RNAseq experiment on neutrophils from healthy volunteers was performed. In addition, neutrophils were stimulated with immobilized IC for 6 h and compared to unstimulated neutrophils to reveal any transcriptional changes. Fig. 1A shows the best-described CDK-cyclin complexes involved in the cell cycle and the transcriptional cycle irrespective of whether their expression changes significantly upon IC stimulation or not, whereas Fig. 1B only shows significantly altered CDKs, cyclins, and selected regulators after IC stimulation. From the 20 CDKs that are overall expressed in humans, we found 14 CDK transcripts (considering both paralogues of *CDK11*) expressed in unstimulated neutrophils and six significantly changed by IC stimulation. While *CDK13* and *CDK19* expression was downregulated after 6 h, *CDK2*, *CDK4*, *CDK7*, and *CDK12* expression was upregulated. Regarding cyclins, 16 out of 29 genes were expressed. On the one hand, the CDKs with best established roles in direct regulation of the cell cycle, namely *CDK1*, *CDK2*, *CDK4*, and *CDK6*, were all detected by RNAseq except *CDK1*. The gene expression of *CDK2* and *CDK4* was induced upon IC stimulation. In addition, their partner cyclins were also detected except cyclin E (*CCNE*), which binds to CDK2 during late G1 phase to promote G1-S transition [90, 91]. Cyclin A (*CCNA*) was unchanged after IC stimulation, whereas cyclin B (*CCNB*) and D (*CCND*) showed an increased expression. On the other hand, expression of the transcriptional kinases *CDK7*, *CDK9*, *CDK12*, *CDK13* and *CDK19* was observed, but not of *CDK18*. Their cyclin binding-partners were all detected with cyclin H (*CCNH*) and T (*CCNT*) elevated upon IC treatment. In addition, the gene expression of other CDK-regulating proteins like RB1, EP300 or MED12/13 was significantly elevated after IC stimulation (Fig. 1B). Furthermore, GO pathway analysis of all genes significantly changed by IC stimulation revealed an activation of granulocyte-specific pathways and a suppression of metabolic pathways (Fig. 1C).

**Fig. 1:**
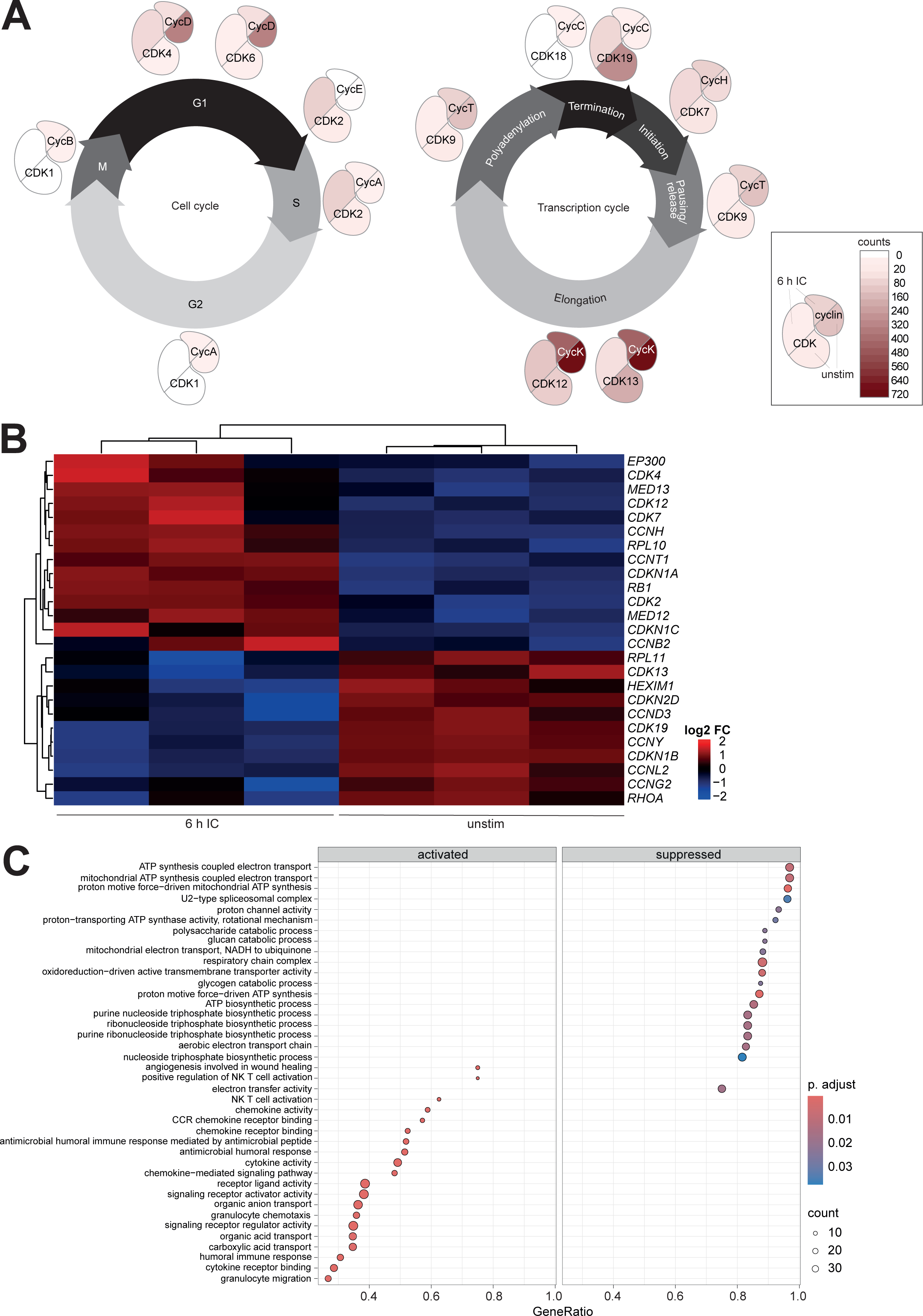
Cell-cycle and transcriptional *CDKs* are expressed in human neutrophils and IC-stimulation partially alters their expression. (**A**) Human neutrophils were purified and unstimulated cells (unstim) plus cells stimulated for 6 h with COL7 anti-COL7 IgG1-IC (6 h IC) were analyzed by bulk RNAseq. Expression in unstimulated and stimulated neutrophils was compared for the individual *CDKs* on the basis of raw counts. **(B)** Significantly differentially expressed genes among CDKs, cyclins, and selected regulators were determined and log2 fold changes (log2 FC) are displayed for the individual samples. **(C)** Top 20 GO activated and suppressed pathways after gene set enrichment analysis of genes significantly differentially expressed upon IC stimulation. Adjusted p value (p. adjust) < 0.1. Gene ratio equals the ratio of differentially expressed genes in the given GO term, count refers to the absolute number. n = 3.

### qRT-PCR confirms the expression of 14 CDKs from RNAseq in human PMNs

To validate the results from RNAseq, an independent qRT-PCR experiment was performed on unstimulated and IC-stimulated human PMNs. Data are provided in Additional file 1. All tested *CDKs* were detected, although *CDK1*, *CDK3*, *CDK5*, *CDK6*, *CDK8*, *CDK11A*, and *CDC7* were expressed at a relatively low level (2^-ΔΔCt^ < 0.1) under both conditions. The expression of *CDK2* and *CDK4* was increased more than threefold or sixfold, respectively, after IC stimulation, while *CDK9* expression decreased by half and *CDK19* expression by approximately 70 %. *CDK3* and *CDK11B* showed a trend to be reduced (both p = 0.0556), but all other *CDKs* remained unaffected by IC stimulation (Fig. 2A). The results support the RNAseq data which also indicate an IC-mediated induction of *CDK2* and *CDK4* transcription, but a reduction of *CDK9* and *CDK19* transcription (see section above). For comparison with proliferating cells, B and CD4^+^ T cells were also analyzed in qRT-PCR (Fig. 2B). Strikingly, *CDK1*, *CDK2*, *CDK4*, *CDK5*, *CDK6*, *CDK8*, and *CDC7* were expressed at higher levels in B and/or T cells, whereas *CDK4*, *CDK9*, *CDK11B*, *CDK14*, and *CDK19* were increased in PMNs.

**Fig. 2:**
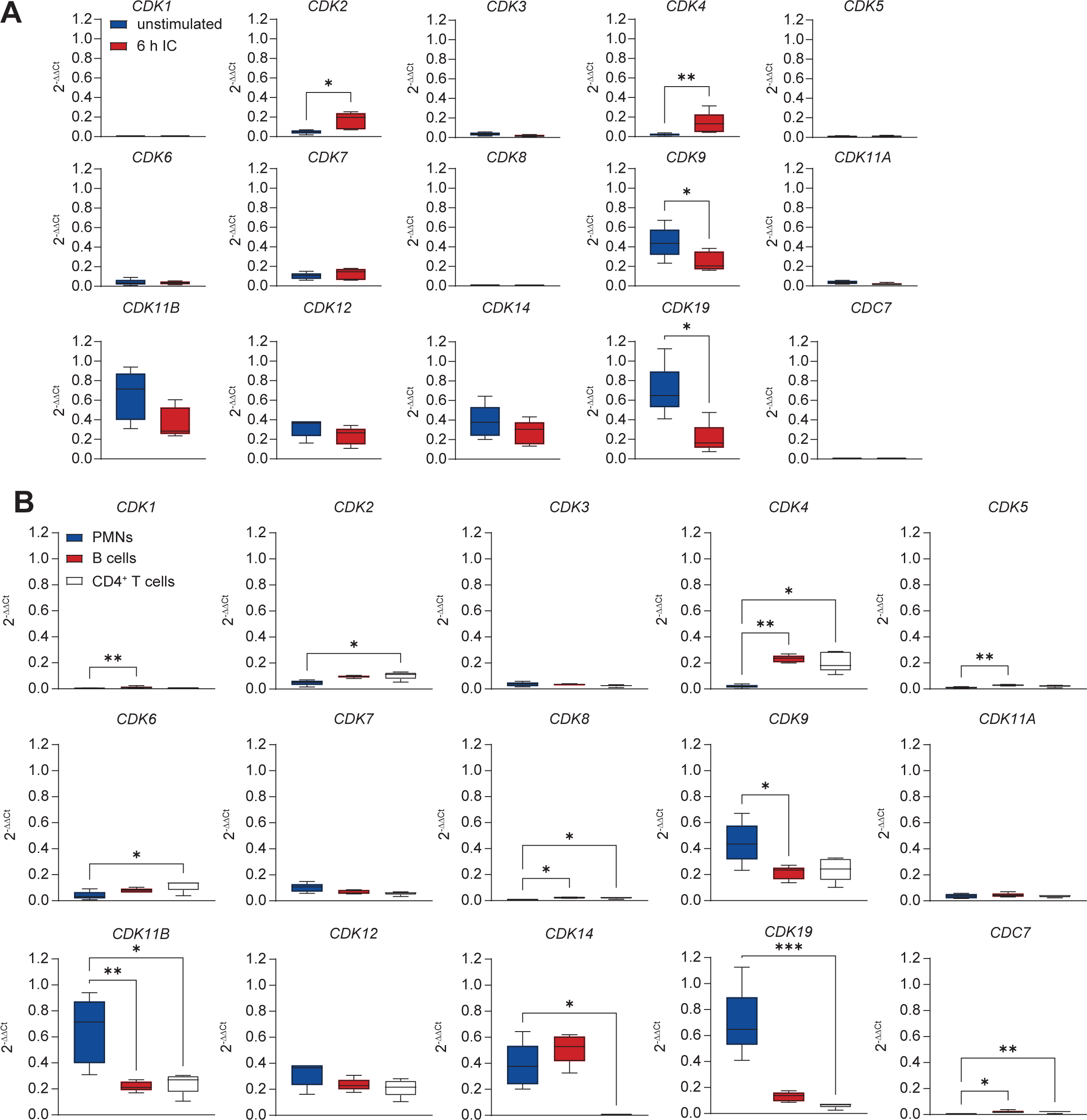
*CDK2* and *CDK4* expression are induced upon IC stimulation in human PMNs. **(A)** Human PMNs were isolated, stimulated for 6 h with COL7 anti-COL7 IgG1-IC and subjected to qRT-PCR detecting 14 distinct CDKs and CDC7 using the the 2^-ΔΔCt^ method. **(B)** In parallel, B cells and CD4^+^ T cells were purified and analyzed in the same way for comparison with unstimulated PMNs. N = 5. Mann-Whitney test **(A)** or Kruskal-Wallis test with Dunn’s multiple comparisons test **(B)**. * p ≤ 0.05. ** p ≤ 0.01. *** p ≤ 0.001.

### Pharmacological inhibition of multiple specific CDKs partially protects *in vitro* from neutrophil effector functions in a dose-dependent manner

To functionally evaluate the role of CDKs in neutrophil kinase signaling, we selected CDKi based on an extensive literature review and tested them in several functional neutrophil assays, covering multiple steps from neutrophil activation to cell death. Therefore, COL7-anti-COL7-IgG1 IC were immobilized on plates, incubated with PMNs from healthy donors and the respective experiment was performed. The surface expression of CD62L, an L-selectin which is shed upon neutrophil activation [92, 93], was analyzed first. Data are provided in Additional file 2. The CDK9i MC180925, CDK7i THZ1 2HCl (Fig. 3), and CDK1i Ro 3306 (Fig. S1) significantly reduced CD62L shedding, while the other inhibitors tested did not (Fig. 3, S1). CDK7 inhibition with THZ2 instead significantly reduced CD18 expression (Fig. 3), which is a β2-integrin responsible for neutrophil adhesion [94]. Similar effects were observed for Ro 3306 and THZ1 2HCl (Fig. S2). Subsequently, adhesion measurements demonstrate a significant reduction only for THZ2 (Fig. 3, S3). Furthermore, THZ2, MC180295, OTS964, Ro 3306, BMS-265246, K03861, palbociclib HCl, THZ1 2HCl, and BAY1251152 inhibited ROS release (Fig. 3, S4). Regarding apoptosis, a significant increase was measured for THZ2, OTS964, and BSJ-4-116 (Fig. 3, S5). For BMS-265246 and AS2863619, statistical significance was reached, however, the values did not exceed those of the negative controls (Fig. S5). Although viability was significantly lower, especially at 10 µM, more than 99 % of the cells remained viable in all cases (Fig. 3, S6). Taken together, CDK inhibition impairs various aspects of IgG-induced neutrophil activation, with ROS release being significantly reduced across multiple inhibitors at doses not substantially affecting viability. Thus, the inhibitory potential of THZ2, MC180295, and OTS964, in particular, is highlighted.

**Fig. 3:**
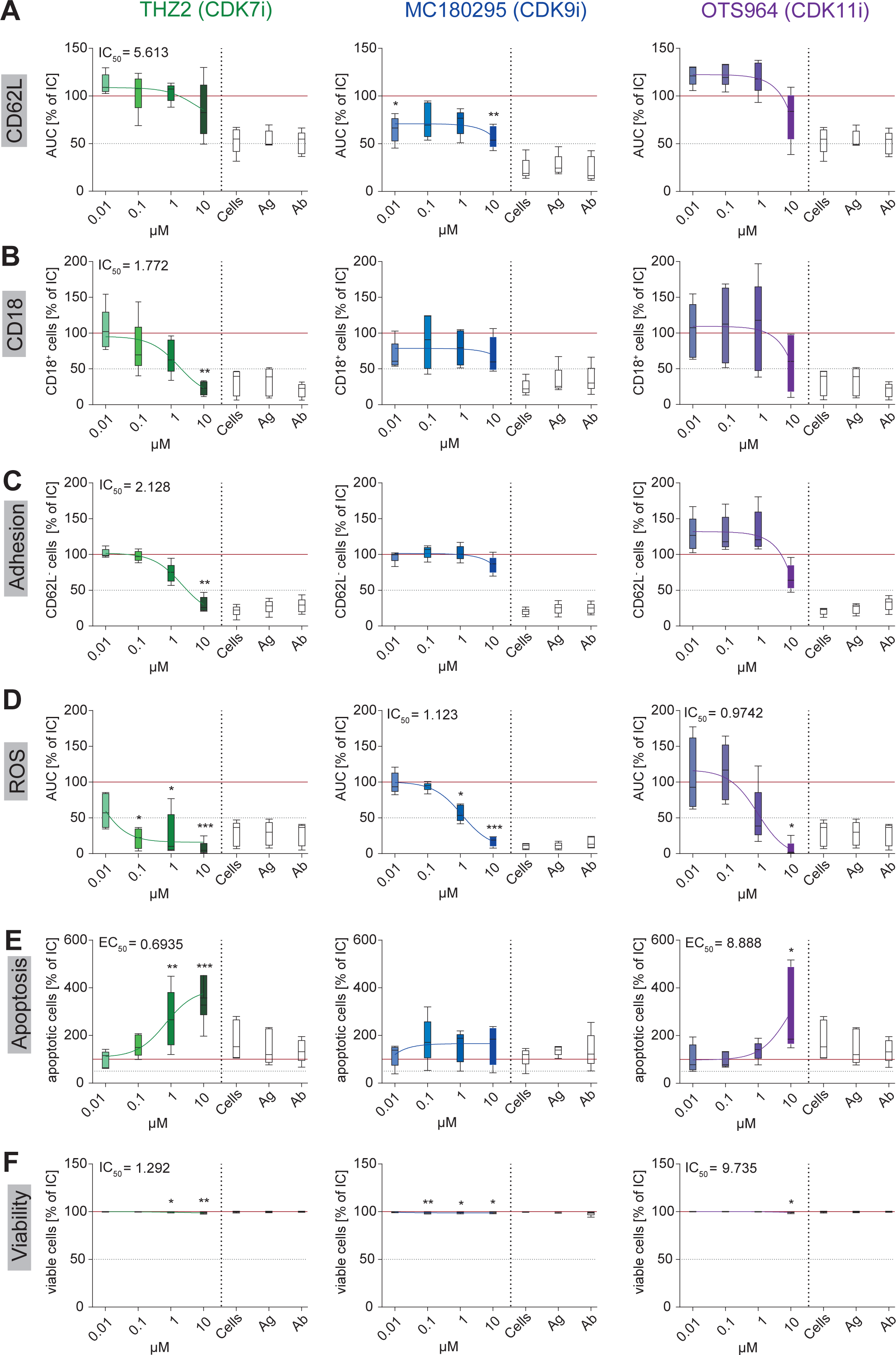
CDK7, CDK9, and CDK11 inhibition attenuate neutrophil effector functions and trigger apoptosis. Different neutrophil functions and characteristics were analyzed after COL7 anti-COL7 IgG1-IC stimulation of human PMNs for 2 h and inhibition with three CDKi: **(A)** CD62L shedding, **(B)** CD18 expression, **(C)** adhesion, **(D)** ROS release, **(E)** apoptosis, and **(F)** viability. The results were normalized to the positive control stimulated with IgG1-IC but without inhibitor indicated by the red line. The dose-response curves were calculated and the IC_50_/EC_50_ values are indicated if in the tested concentration range. Cells only, antigen only (Ag), and antibody only (Ab) served as negative controls. Kruskal-Wallis test with Dunn’s multiple comparisons test was performed. * p ≤ 0.05. ** p ≤ 0.01. *** p ≤ 0.001. For apoptosis experiment using THZ2 n = 6, rest n = 5. AUC, area under the curve.

### Both systemic and topical treatment with CDK7, CDK9 or CDK11 inhibitors reduces disease severity in the local antibody-transfer induced EBA model

Following the successful *in vitro* validation of the selected CDKi, a local antibody-transfer induced EBA model was performed to test these compounds as systemic treatment (Fig. 4A). Data are provided in Additional file 3. CDK9 inhibition by MC180295 completely prevented disease induction, which is exemplified by representative images showing the ameliorated clinical presentation of the mice and the reduced inflammation in the H&E staining despite IgG and C3 deposition along the dermal-epidermal junction (Fig. 4B). The AESA was significantly reduced starting at day 1 and increasing further until day 3, which is also reflected by the H&E score (Fig. 4C). CDK7 inhibition by both THZ2 (Fig. 4D) and THZ1 2HCl (Fig. 4E) reduced the AESA, but only THZ2 treatment significantly decreased the H&E score. Systemic CDK11 inhibition by OTS964 treatment also ameliorated the disease severity, but the H&E score did not change significantly between the treated and the control mice (Fig. 4F). However, treatment with Ro 3306 and palbociclib HCl had no significant effect (Fig. S7).

**Fig. 4:**
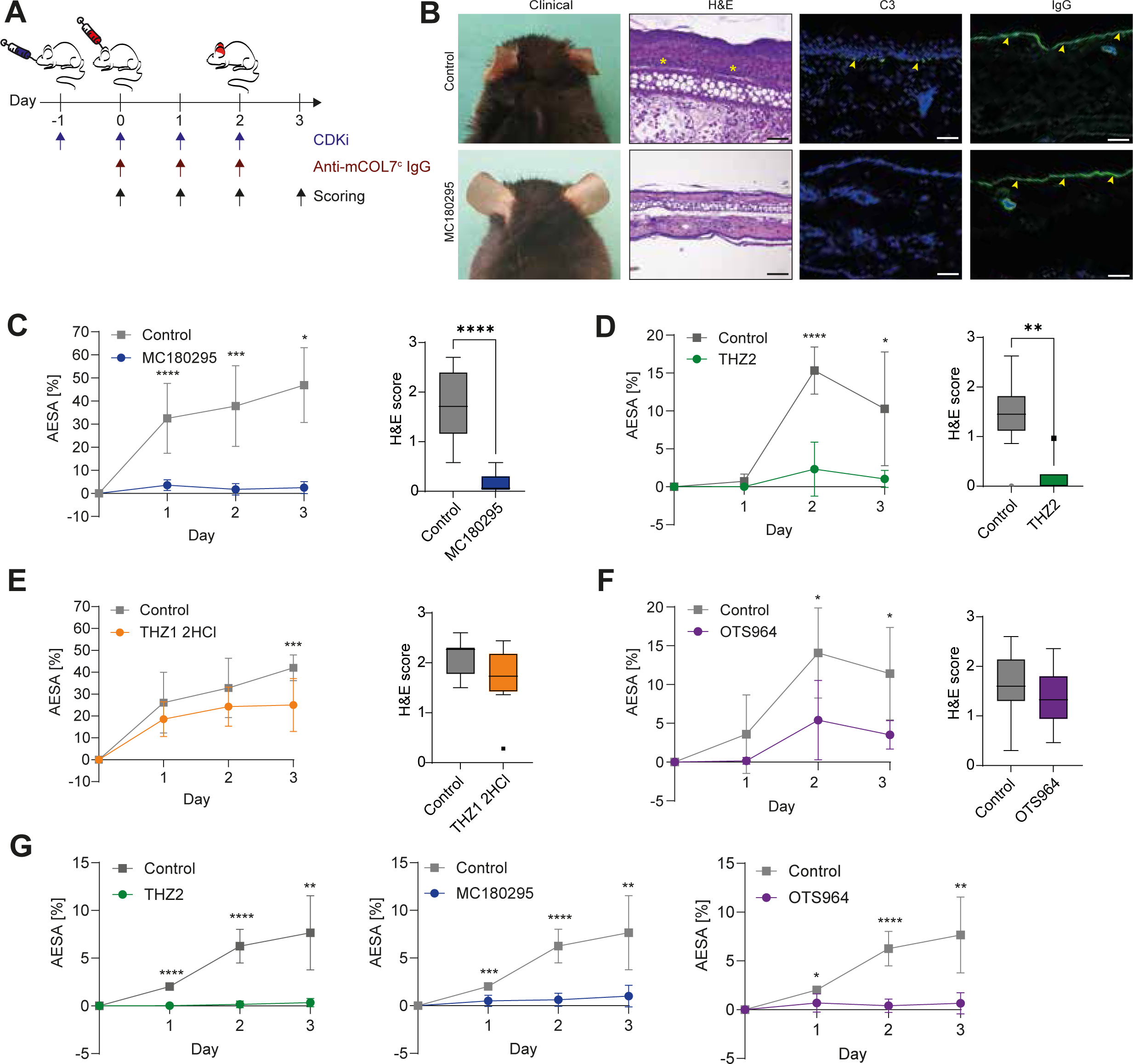
Systemic and topical CDK7i, CDK9i, and CDK11i treatment reduces disease severity in the local EBA. **(A)** EBA was induced by injection of anti-mCOL7^c^ IgG and the mice were treated with CDKi either systemically or topically. **(B)** Exemplary clinical and histological images are shown with the scale bars corresponding to 100 µM. Asterisks indicate cell infiltrates and arrows indicate binding of C3/IgG to the dermal-epidermal junction. **(C)** The affected ear surface area (AESA) and the H&E score were evaluated in the systemic treatment group for MC180295, **(D)** THZ2, **(E)** THZ1 2HCl, and **(F)** OTS964. **(G)** For the topical model, AESA was analyzed as well. Two-way ANOVA with Šidák’s multiple comparisons test was performed for AESA evaluation, except for the systemic treatment with THZ2 and MC180295 a mixed-effects analysis with Šidák’s multiple comparisons test was conducted. For H&E scores, Mann-Whitney test was carried out. * p ≤ 0.05. ** p ≤ 0.01. **** p ≤ 0.0001. n = 8.

The most promising inhibitors from the systemic treatment in local antibody-transfer induced EBA were selected for topical treatment in the same model. THZ2 was chosen over THZ1 HCl since both inhibit CDK7, but THZ2 showed better results *in vitro* and almost completely protected from disease development *in vivo*. All three inhibitors tested prevented disease induction when applied topically (Fig. 4G).

### CDK9 inhibition with MC180295 is beneficial in neutrophil-dependent STA, but not in neutrophil-independent ITP

The CDKi that showed the most prominent effect in the local antibody-transfer induced EBA model, MC180295, was validated in two other IC-driven models that differ regarding the role of neutrophils in their respective pathogenesis, namely STA and ITP. Data are provided in Additional file 4. In STA (Fig. 5A), systemic treatment with MC180295 significantly reduced the disease score of paw swelling (Fig. 5B). This was reflected in histological analysis showing reduced proteoglycan degradation and thus cartilage damage as compared to control mice upon staining with Safranin-O. H&E staining revealed pronounced immune cell infiltration in the controls, but not or to a lesser extent in the treated mice (Fig. 5C). Out of 13 cytokines tested in the serum, two were significantly changed on the final day between the two groups. Both IL-6 and IL-10 were reduced on day 7 in the MC180295-treated group compared to the control group. All other measured cytokine levels were not significantly altered over the course of the observation period of 7 days (Supplemental table 2).

**Fig. 5:**
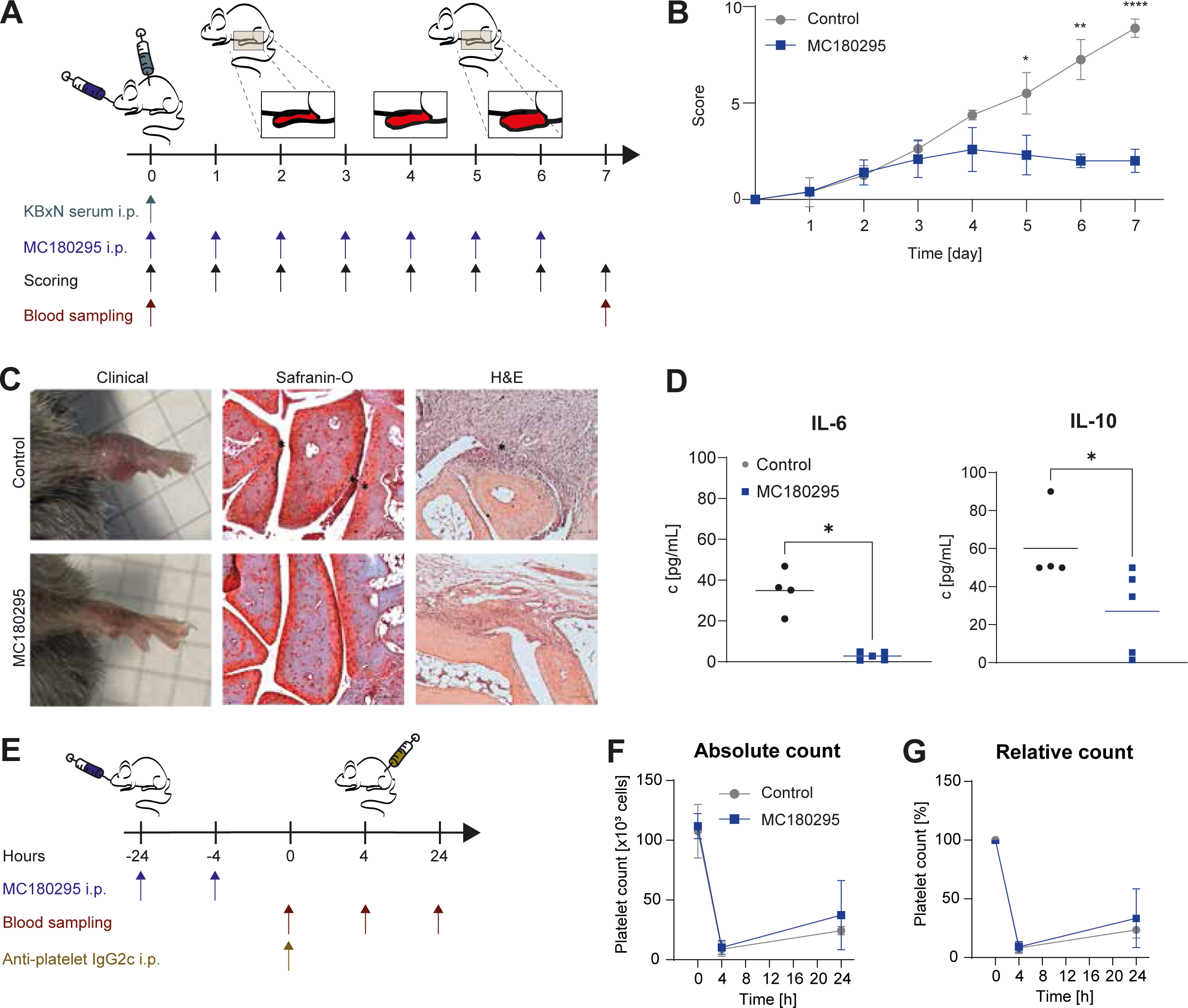
CDK9 inhibition with MC180295 improves disease severity in STA, but not in ITP. **(A)** To induce STA, mice were injected with KBxN serum and treated with MC180295 for seven days. **(B)** The course of the disease score is depicted and **(C)** reflected by the clinical presentation of the paws and the histological stainings. The scale bars represent 100 µm. Asterisks indicate proteoglycan staining (Safranin-O) or cell infiltration (H&E). **(D)** Cytokine levels were measured in serum and significantly changed cytokines are shown. **(E)** ITP was induced by anti-platelet IgG2c-induced platelet depletion and MC180295 was applied at 24 h and 4 h before depletion. **(F)** Absolute and **(G)** relative platelet counts were monitored for up to 24 h and values normalized to 100 % at 0 h are indicated. For arthritis score and platelet counts, two-way ANOVA with Šidák’s multiple comparisons test was performed. For cytokine levels, Mann-Whitey test was conducted. * p ≤ 0.05. ** p ≤ 0.01. **** p ≤ 0.0001. **(A-D)** n = 5 (treatment) and n = 4 (control). **(E-G)** n = 4.

**Supplemental table 2.** Descriptive statistics for all cytokines analysed in STA at day 0 and day 7.

| day 0 |  |  |  | day 7 |  |  |
| --- | --- | --- | --- | --- | --- | --- |
| median ± SEM |  | p value |  | median ± SEM |  | p value |
| cyto- | treated | control | treated vs. | treated | control | treated vs. |
| kine |  |  | control |  |  | control |
| **GM-** | 0.564 ± | 0 ± 0 | 0.4444 | 1.036 ± | 1.465 ± | 0.5556 |
| **CSF** | 0.3987 |  |  | 0.6401 | 0.7801 |  |
| **IL-10** | 13.99 ± | 16.63 ± | 0.6825 | 27.06 ± | 60.15 ± | 0.0317 |
|  | 8.837 | 16.63 |  | 9.973 | 9.976 |  |
| **IL-** | 1.58 ± | 0.69 ± | 0.5714 | 1.546 ± | 3.53 ± | 0.3095 |
| **12p70** | 0.8095 | 0.3984 |  | 1.029 | 1.436 |  |
| **IL-17A** | 1.144 ± | 0.44 ± | 0.5556 | 1.53 ± | 1.47 ± | 0.9048 |
|  | 0.5896 | 0.2 |  | 0.4803 | 0.3853 |  |
| **IL-1α** | 190 ± | 218.5 ± | 0.5556 | 15.69 ± | 14.89 ± | 0.7302 |
|  | 41.09 | 33.07 |  | 4.246 | 5.507 |  |
| **IL-1β** | 7.224 ± | 4.81 ± | > 0.9999 | 3.076 ± | 7.043 ± | 0.1032 |
|  | 4.069 | 1.707 |  | 1.342 | 1.508 |  |
| **IL-23** | 29.37 ± | 15.08 ± | 0.4365 | 43.63 ± | 25.55 ± | 0.5476 |
|  | 11.81 | 9.157 |  | 17.1 | 6.944 |  |
| **IL-27** | 124.5 ± | 133.7 ± | 0.7302 | 116.6 ± | 111.8 ± | 0.375 |
|  | 32.39 | 85.19 |  | 41.59 | 18.37 |  |
| **IL-6** | 4.162 ± | 2.088 ± | 0.1746 | 2.824 ± | 34.84 ± | 0.0159 |
|  | 1.272 | 1.033 |  | 0.8739 | 5.281 |  |
| **IFN-β** | 78.04 ± | 37.03 ± | 0.4127 | 10.04 ± | 29.55 ± | 0.0952 |
|  | 20.83 | 27.01 |  | 6.495 | 7.633 |  |
| **IFN-γ** | 3.578 ± | 2.833 ± | 0.7937 | 2.968 ± | 5.545 ± | 0.127 |
|  | 1.287 | 0.9399 |  | 0.6265 | 0.9739 |  |
| **MCP-1** | 12.09 ± | 7.603 ± | 0.2857 | 9.492 ± | 16.65 ± | 0.3175 |
|  | 2.878 | 3.671 |  | 5.536 | 4.481 |  |
| **TNF-α** | 13.24 ± | 11.38 ± | 0.4444 | 10.42 ± | 22.59 ± | 0.0952 |
|  | 2.719 | 1.989 |  | 3.304 | 6.41 |  |

In contrast to STA, ITP (Fig. 5E), where neutrophils are not central to disease pathogenesis, prophylactic treatment with MC180295 did not show any effects on platelet count 24 h after platelet depletion (Fig. 5G), demonstrating the variable effect of CDKi in the different disease models.

## Discussion

IC-induced FcγR-mediated kinase signaling plays a major role in intracellular signal transduction by neutrophils under pathological conditions, e.g., in autoimmune diseases such as EBA or RA. However, the role of CDKs in neutrophils is largely unknown. A small number of studies have reported the expression of CDKs on protein level and suggested functional relevance in these cells [22, 28, 95]. To the best of our knowledge, we show in the present study for the first time a comprehensive, functional analysis of IC-triggered FcγR-dependent kinase signaling using a variety of selective CDKi. Fig. 6 summarizes our main findings.

**Fig. 6:**
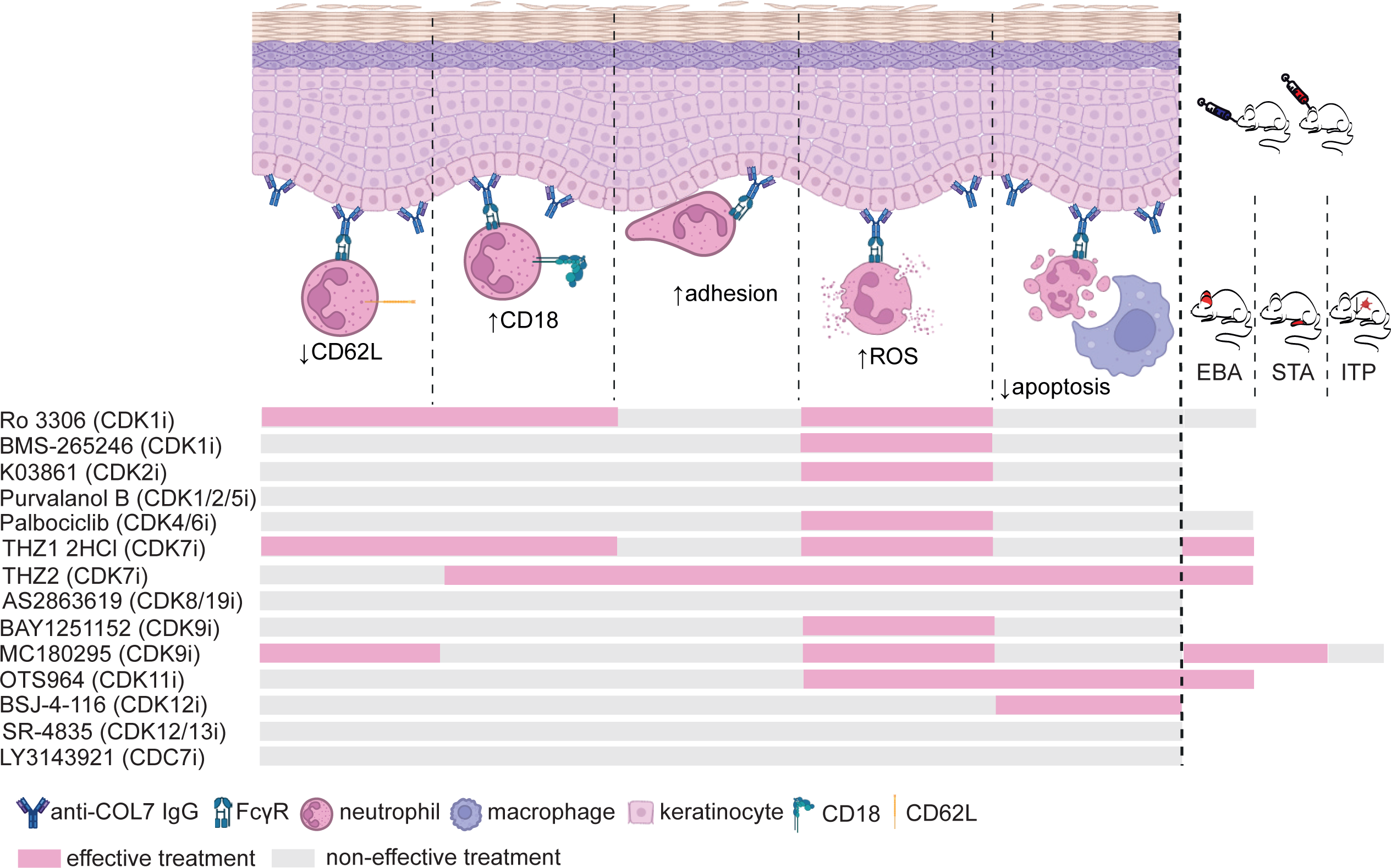
Successful inhibition of neutrophil functions and experimental autoimmune models using CDKi. Out of 20 CDKs that are described in humans, 14 were detected in our gene expression analysis. After choosing selective CDKi, different *in vitro* neutrophil functional assays were performed. The number of efficient inhibitors is indicated. Subsequently, all CDKi that inhibited ROS release were tested as systemic treatment in the local antibody transfer-induced EBA model, except for BAY1251152 to avoid redundancy. Three CDKi reduced the disease severity and were thus applied topically in the same model. Additionally, the most promising CDKi was validated in STA and ITP. Created in https://BioRender.com.

Interestingly, we identified multiple CDKs as expressed at RNA level and found that their transcription may change in IC-stimulated neutrophils. The expression of the two traditional cell-cycle CDKs *CDK2* and *CDK4* increased, whereas the expression of the two transcriptional CDKs *CDK9* and *CDK19* decreased. Concerning CDK9, it has been shown that inhibition leads to the reactivation of epigenetically silenced genes, e.g. *SYNE1*, *GLDC*, *MUC20*, *ANPEP*, *SUSD4*, and *COLEC11* [64]. All of these genes have been linked to the immune system [96], e.g. NETosis [97–99], apoptosis [100], complement inhibition [101], or neutrophil degranulation [102]. Therefore, CDK9 inhibition might further influence these processes.

The trends found in RNAseq are clearly supported by our qRT-PCR analysis in an independent dataset. Furthermore, the qRT-PCR data highlight the elevated expression of *CDK9*, *11B* and *19* in neutrophils compared to B and CD4^+^ T cells, making them especially promising targets in neutrophil-dependent diseases. The tendencies concerning gene expression levels in unstimulated cells found in our RNAseq and qRT-PCR analyses are in line with publicly available RNAseq data of unstimulated neutrophils from six healthy donors sorted by flow cytometry [88, Human Protein Atlas proteinatlas.org]. Several CDKs like CDK2, CDK4, and CDK11 were also detected in a kinase activity measurement of IC-stimulated PMNs [74].

Since CDKs only become fully active after binding to their cyclin binding partners, we also examined cyclins in our RNAseq analysis. Most prominently, we found both CDK7 and its binding partner cyclin H significantly increased upon IC activation. Strikingly, *MNAT1* (MAT1 RNA) was not detected, suggesting the requirement of a phosphorylation-dependent activation of CDK7 [104, 105].

Furthermore, several regulators of CDKs were differentially expressed upon IC stimulation in human neutrophils, hence adding another layer of complexity to the regulation of CDKs in autoimmune disease. Both *MED12* and *MED13* were transcribed more, possibly promoting the formation of a transcriptional pre-initiator complex after association with CDK8/19 and cyclin C [106]. Moreover, regulators of chromatin remodeling required for transcriptional activation like *RB1* [107] and *EP300* [108, 109] were detected at higher RNA levels in stimulated cells.

In addition to the analysis focusing on CDKs and their regulators, we performed a GO pathway analysis for the enriched gene sets from unstimulated versus IC-stimulated neutrophils. Interestingly, our pathway analysis revealed that many metabolic pathways including ATP synthesis or catabolic processes are suppressed, while immune pathways such as granulocyte migration, chemokine and cytokine pathways were activated.

Thus, we hypothesized that targeting CDKs by selective small-molecule inhibitors might reduce neutrophil effector functions in IC-stimulated neutrophils *in vitro* and consequently disease severity in IC-driven neutrophil-dependent models of autoimmune disease *in vivo*. Although there are multiple studies published on CDK inhibition in different contexts, they rarely focus on neutrophils and utilise relatively unselective CDKi. However, pan-CDK inhibition is often associated with severe side effects [110]. Here, we therefore chose more selective CDK inhibitors to investigate their role on neutrophil functions upon IC activation.

Hence, we analyzed several functions that are critical for neutrophil recruitment as well as apoptosis. As multiple studies suggest that ROS release might be a predictor of *in vivo* efficacy in experimental EBA [74, 75, 79, 111], all CDKi that significantly reduced ROS release were tested as systemic treatment in the local antibody transfer-induced EBA model (Ro 3306, palbociclib HCl, THZ1 2HCl, THZ2, OTS964). Moreover, inhibition of CDK1, CDK2, CDK4/6, CDK7, CDK9, or CDK12 influenced other neutrophil functions such as expression of the surface activation markers CD18 and CD62L, and adhesion, highlighting a role for CDKs in neutrophil recruitment and activation. Viability was only slightly influenced, with values still in the range of the negative controls. However, some inhibitors (THZ2, OTS964, BSJ-4-116) increased apoptosis, which is a highly controlled form of cell death leaving the cell membrane intact [25]. In line with our data, a role in regulating apoptosis has already been suggested for CDK7 in neutrophils [22], and for CDK11 in Chinese Hamster Ovary cells and T-cell lines [112]. Concerning neutrophils, this may protect the tissue from toxic substances like ROS which are released during alternative forms of cell death and has been described as a potential mechanism of action of CDKi [95, 113]. Indeed, THZ2 and OTS964 increased apoptosis while reducing ROS. However, other inhibitors reduced ROS release without affecting apoptosis (Ro 3306, K03861, palbociclib HCl, THZ1 2HCl, BAY1251152), implicating the involvement of different mechanisms or signaling pathways. Interestingly, CDK9 inhibition by MC180295 or BAY1251152 did not significantly influence apoptosis, whereas CDK9 inhibition by promiscuous inhibitors such as flavopiridol or R-roscovitine resulted in neutrophil apoptosis induction [22, 30]. Thus, this indicated an inhibitor-specific, probably synergistic effect caused by targeting several CDKs at once.

Subsequently, the most promising CDKi were applied systemically in the local antibody-transfer induced EBA model. Here, inhibition of CDK7 by THZ1 2HCl or THZ2, CDK9 by MC180295, or CDK11 by OTS964 were found to significantly reduce the disease severity. Remarkably, CDK7, CDK9, and CDK11 are all described as transcriptional kinases [3]. This is in line with our expectation considering that neutrophils are non-proliferating cells. Inhibition of CDK1 with Ro 3306 or CDK4/6 with palbociclib HCl, in contrast, did not reduce the AESA. Although palbociclib HCl showed promising effects *in vitro* and is licensed as treatment for breast cancer, where it improves the neutrophil-to-lymphocyte ratio in combination with the aromatase inhibitor letrozole [114], other data indicate an opposite role for CDK6 in regulating NET formation as CDK6^-/-^ mice failed to produce NETs upon PMA stimulation [28]. Additionally, CDK6^-/-^ mice exhibited a higher amount of primary granule proteins, suggesting CDK6 as a regulator of neutrophil granularity [115]. These data might explain the outcome seen in EBA.

The most efficient CDKi, MC180295, was next administered in two other models to verify the neutrophil-dependent effects: STA, which is neutrophil- and macrophage-dependent, and ITP, which is mostly neutrophil-independent. In STA, MC180295 significantly reduced disease severity and levels of several inflammatory cytokines. These findings complement a study describing protective effects on human chondrocytes and cartilage explants by CDK9 inhibition with flavopiridol or small interfering RNA. Less pro-inflammatory cytokines are released, preventing cartilage degradation [32]. Of note, recent data show the induction of neutrophil reverse migration from the inflammation site back to the circulation upon CDK9 inhibition in zebrafish larvae, hence contributing to the resolution of inflammation [116].

Nevertheless, compared to the almost complete protection from disease development observed in the local antibody transfer-induced EBA model, the MC180295-treated STA mice showed a residual mean arthritis score of 2 on the final day, representing still 22 % of that observed in the control mice, in contrast to only 5 % in the EBA model. This finding might reflect the different functional relevance of neutrophils in the two distinct models. While EBA is described to be primarily neutrophil-driven [38], neutrophils also play a critical [45], yet not exclusive, role in STA [47]. In the latter model, the function of macrophages, NK cells, mast cells, and synovial fibroblasts during the effector phase of the disease is also well established [44] and might explain the differences in efficacy seen on both models. A study conducted with LPS-stimulated human monocyte-derived macrophages demonstrated a reduction of IL-6 and TNF after treatment with the more promiscuous CDK9i AT7519 or roscovitine, supporting a role for macrophages in the resolution of inflammation [117]. In addition, administration of the CDK9i PC585 and PC579 in murine arthritis models led to the downregulation of the anti-apoptotic protein MCl-1, thus favoring apoptosis [118]. As delayed apoptosis is a feature of RA [46], CDK inhibition might be a reasonable approach in this context.

Conversely, no effects on platelet depletion or recovery were observed in ITP. Contrasting with EBA and STA, neutrophils are considered dispensable for the effector phase of ITP. Rather other immune cells like tissue-resident macrophages are involved [51–53]. This highlights the role for CDK9 in IC-induced neutrophil-dependent diseases like EBA and STA.

Regarding potential limitations of this study, only one concentration of each inhibitor was used *in vivo*. Thus, any effects which might be seen at different concentrations remain unclear. However, these concentrations had successfully been used in mouse models before and were hence deemed safe to apply. For example, MC180295 demonstrated an anti-tumoral effect in NOD.Cg-*Prkdc^scid^ Il2rg^tm1Wjl^*/SzJ mice lacking mature B and T cells [64] and palbociclib is already in clinical use for the treatment of metastatic breast cancer [119]. Furthermore, our *in vitro* data indicate a mechanistic effect of CDKi rather than a mere cytotoxic effect.

Additionally, only one inhibitor per target (except for CDK7 since both inhibitors were equally promising) was tested. We are aware that other inhibitors with different selectivity or bioavailability might be more efficient, nevertheless the use of further inhibitors would have exceeded the scope of this explorative study. The way is now paved for further studies grounding on this work.

Instead of pharmacological inhibition, genetic knockouts could be a more precise alternative to address our research questions. Unfortunately, genetic deletions of many CDKs are embryonically lethal, with CDK1 being indispensable for early embryogenesis [120, 121]. CDK4- and CDK6-deficient mice display problems in embryonic development [122, 123] similar to CDK8^-/-^ [124], CDK11^-/-^ [125] or CDK12^-/-^ mice [126]. Therefore, we consider pharmacological inhibition to be a practicable and clinically relevant alternative, though future studies with neutrophil-specific inducible depletion may be informative.

## Conclusions

Taken together, we have demonstrated the functional relevance of CDKs including CDK7, CDK9, and CDK11 in neutrophil IC-induced kinase signaling both *in vitro* and *in vivo*. Our findings contribute to the exploration of potential novel treatment options for autoimmune diseases such as PD or RA.

## List of abbreviations

AUC: area under the curve
BSA: bovine serum albumin
CAK: CDK-activating kinase
CDK: cyclin-dependent kinase
CDKi: small-molecule CDK inhibitor
CKI: intrinsic CDK inhibitor
COL7: collagen VII
DMSO: dimethyl sulfoxide
EBA: epidermolysis bullosa acquisita
ENA: European Nucleotide Archive
FcγR: fragment crystallisable gamma receptor
GO: Gene Ontology
H&E: hematoxylin and eosin
i.p.: intraperitoneal
IC: immune complex
ITP: immune thrombocytopenia
MACS: magnetic activated cell sorting
NET: neutrophil extracellular trap
PBS: phosphate-buffered saline
PD: pemphigoid disease
PMN: polymorphonuclear granulocyte
qRT-PCR: real-time quantitative reverse transcription polymerase chain reaction
RA: rheumatoid arthritis
RNAseq: bulk RNA sequencing
ROS: reactive oxygen species
RT: room temperature
STA: serum-transfer arthritis

## Declarations

### Consent for publication

Not applicable

### Availability of data and materials

The RNAseq dataset generated and analyzed during the current study is available in the European Nucleotide Archive (ENA) at EMBL-EBI under accession number PRJEB103987 (https://www.ebi.ac.uk/ena/browser/view/PRJEB103987), all other data generated or analyzed during this study are included in this published article and its supplementary information files.

### Author’s contributions

Investigation: MS, LV, APM, DM, SJ, MR, AKS, SMV, YX, HG, JL, writing (original draft): MS, visualization: MS, LV, AKS, formal analysis: CO, NE, AV, MS, AKS, LV, KA, resources: RV, GV, FP, XY, RJL, KTA, conceptualization: RJL, AL, KB, supervision: AL, KB, KTA, software: AV, project administration: AL, KB, writing (review & editing): all.

## Acknowledgements

We thank Kathrin Kalies, Petra Lau, Astrid Fischer, Claudia Kauderer, Alexandra Wobig, Daniela Rieck, Norbert Reiling, Carolin Möller, and Cindy Jensen for their support.

## Figure Legends

**Fig. S1: CDK1i Ro 3306 and CDK7i THZ1 2HCl reduce CD62L shedding in IC-stimulated human PMNs. (A-K)** Human PMNs were stimulated for 2 h with hCOL7 anti-COL7 IgG1-IC and subsequently analyzed for CD62L shedding. Inhibitors targeting specific CDKs were tested in flow cytometry for their ability to minimize CD62L shedding. Based on FSC-SSC gating, alive singlet CD45^+^ cells were considered. The results were normalized to the positive control stimulated with IgG1-IC but without inhibitor indicated by the red line. The dose-response curves were calculated and the IC_50_/EC_50_ values are indicated if in the tested concentration range. Cells only, antigen only (Ag), and antibody only (Ab) served as negative controls. Kruskal-Wallis test with Dunn’s multiple comparisons test was performed. * p ≤ 0.05. ** p ≤ 0.01. *** p ≤ 0.001. n = 5.

**Fig. S2: CDK1i Ro 3306 or CDK7i THZ1 2HCl decrease CD18 expression on IC-activated human PMNs. (A-K)** Following stimulation of human PMNs for 2 h with hCOL7 anti-COL7 IgG1-IC and treatment with CDKi, the cells were analyzed for their CD18 expression in flow cytometry. Based on FSC-SSC gating. alive singlet CD45^+^ cells were considered. The results were normalized to the positive control stimulated with IgG1-IC but without inhibitor indicated by the red line. The dose-response curves were calculated and the IC_50_ values are indicated if in the tested concentration range. Cells only, antigen only (Ag), and antibody only (Ab) served as negative controls. Kruskal-Wallis test with Dunn’s multiple comparisons test was performed. * p ≤ 0.05. ** p ≤ 0.01. n = 5.

**Fig. S3: Adhesion of IC-activated human PMNs is not significantly reduced by the additional CDKi used. (A-K)** Human PMNs were stimulated with hCOL7 anti-COL7 IgG1-IC for 2 h. Impedance-based measurement of adhesion was conducted using inhibitors targeting specific CDKs. The area under the curve was determined and the results were normalized to the positive control stimulated with IgG1-IC but without inhibitor indicated by the red line. The dose-response curves were calculated and the IC_50_/EC_50_ values are indicated if in the tested concentration range. Cells only, antigen only (Ag), and antibody only (Ab) served as negative controls. Kruskal-Wallis test with Dunn’s multiple comparisons test was performed. * p ≤ 0.05. ** p ≤ 0.01. *** p ≤ 0.001. n = 5.

**Fig. S4: Inhibition of CDK1, CDK2, CDK4/6, CDK7, or CDK9 in IC-stimulated human PMNs minimizes ROS release. (A-K)** Activation of human PMNs with hCOL7 anti-COL7 IgG1-IC for 2 h, treatment with selective CDKi and subsequent chemiluminescence-based analysis of ROS release was performed. The area under the curve (AUC) was calculated and the results were normalized to the positive control stimulated with IgG1-IC but without inhibitor indicated by the red line. The dose-response curves were calculated and the IC_50_ values are indicated if in the tested concentration range. Cells only, antigen only (Ag), and antibody only (Ab) served as negative controls. Kruskal-Wallis test with Dunn’s multiple comparisons test was performed. * p ≤ 0.05. ** p ≤ 0.01. *** p ≤ 0.001. n = 5.

**Fig. S5: Apoptosis in IC-stimulated human PMNs is increased by application of CDK1/2i, CDK12i, or CDK8/19i. (A-K)** Human PMNs were activated with hCOL7 anti-COL7 IgG1-IC for 2 h. After using selective CDKi, flow cytometry was performed. Based on FSC-SSC gating, singlet CD45^+^ cells were analyzed. Annexin V^+^ Zombie NIR^-^ cells were considered apoptotic. The results were normalized to the positive control stimulated with IgG1-IC but without inhibitor indicated by the red line. The dose-response curves were calculated and the EC_50_ values are indicated if in the tested concentration range. Cells only, antigen only (Ag), and antibody only (Ab) served as negative controls. Kruskal-Wallis test with Dunn’s multiple comparisons test was performed. * p ≤ 0.05. ** p ≤ 0.01. *** p ≤ 0.001. For apoptosis experiment using Palbociclib HCl, AS2863619, BMS-265246, and SR-4835 n =6, rest n = 5.

**Fig. S6: Viability of IC-treated human PMNs is minorly affected by CDK1, CDK7, and CDK12 inhibition. (A-K)** Human PMNs stimulated with hCOL7 anti-COL7 IgG1-IC and treated with CDKi for 2 h were analyzed for toxic effects in flow cytometry. Based on FSC-SSC gating, singlet CD45^+^ cells were studied. Annexin V^-^ Zombie NIR^-^ cells were considered viable. The results were normalized to the positive control stimulated with IgG1-IC but without inhibitor indicated by the red line. The dose-response curves were calculated and the IC_50_ values are indicated if in the tested concentration range. Cells only, antigen only (Ag), and antibody only (Ab) served as negative controls. Kruskal-Wallis test with Dunn’s multiple comparisons test was performed. * p ≤ 0.05. n = 5.

**Fig. S7: CDK1i Ro 3306 or CDK4/6i palbociclib HCl do not reduce the AESA in local EBA**. **(A-B)** EBA was induced by injection of anti-mCOL7^c^ IgG and the mice were treated systemically with three more CDKi. The affected ear surface area (AESA) was evaluated. Mixed-effects analysis with Šidák’s multiple comparisons test was performed. *** p ≤ 0.001. **(A)** n = 7, **(B)** n = 4.

