## Supplementary figures and images for "CDK7, CDK9, or CDK11 Inhibition Reduces Neutrophil-Driven Inflammation and Tissue Damage in Experimental Autoimmune Models"

### Fig. S1

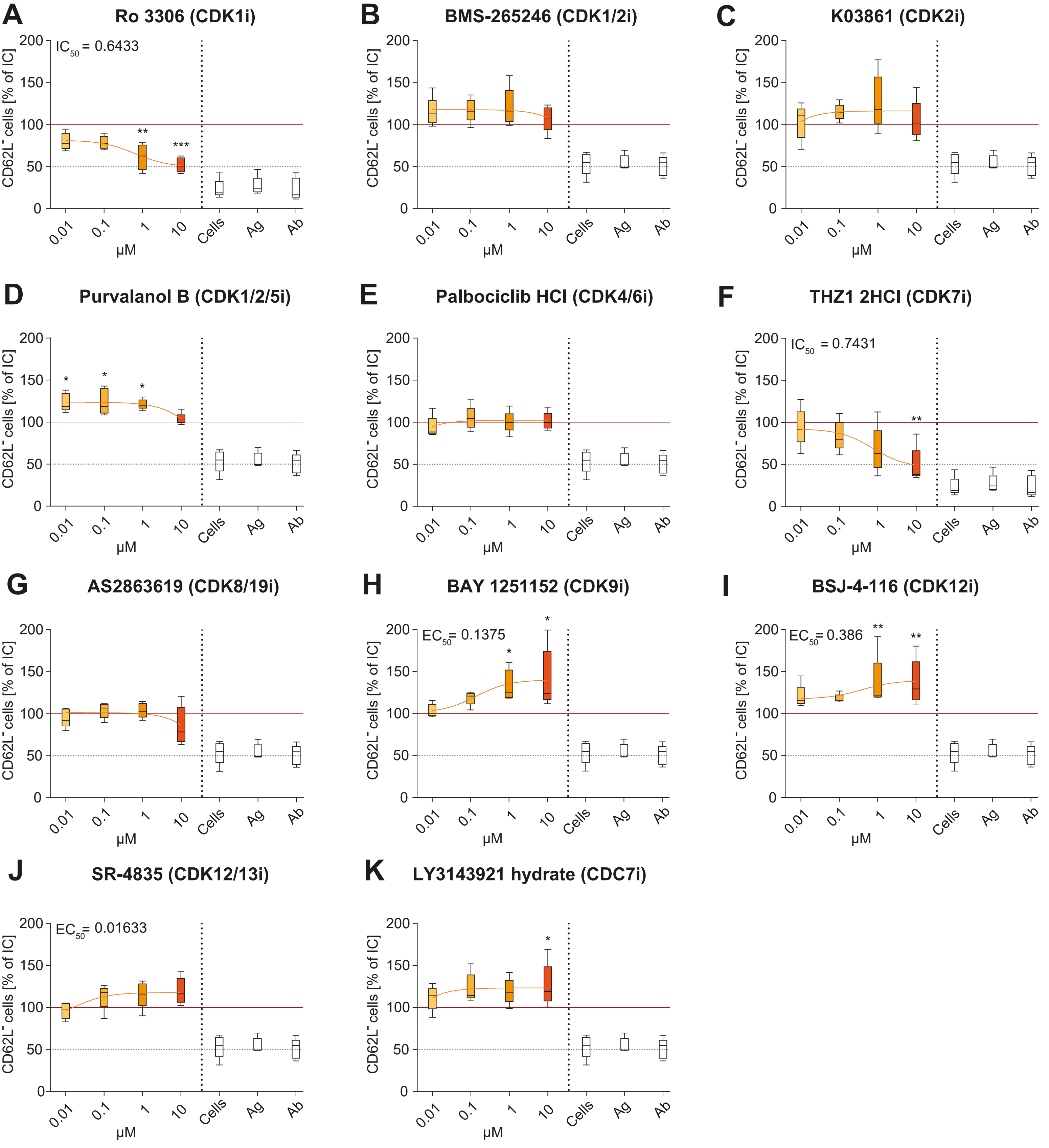

### Fig. S2

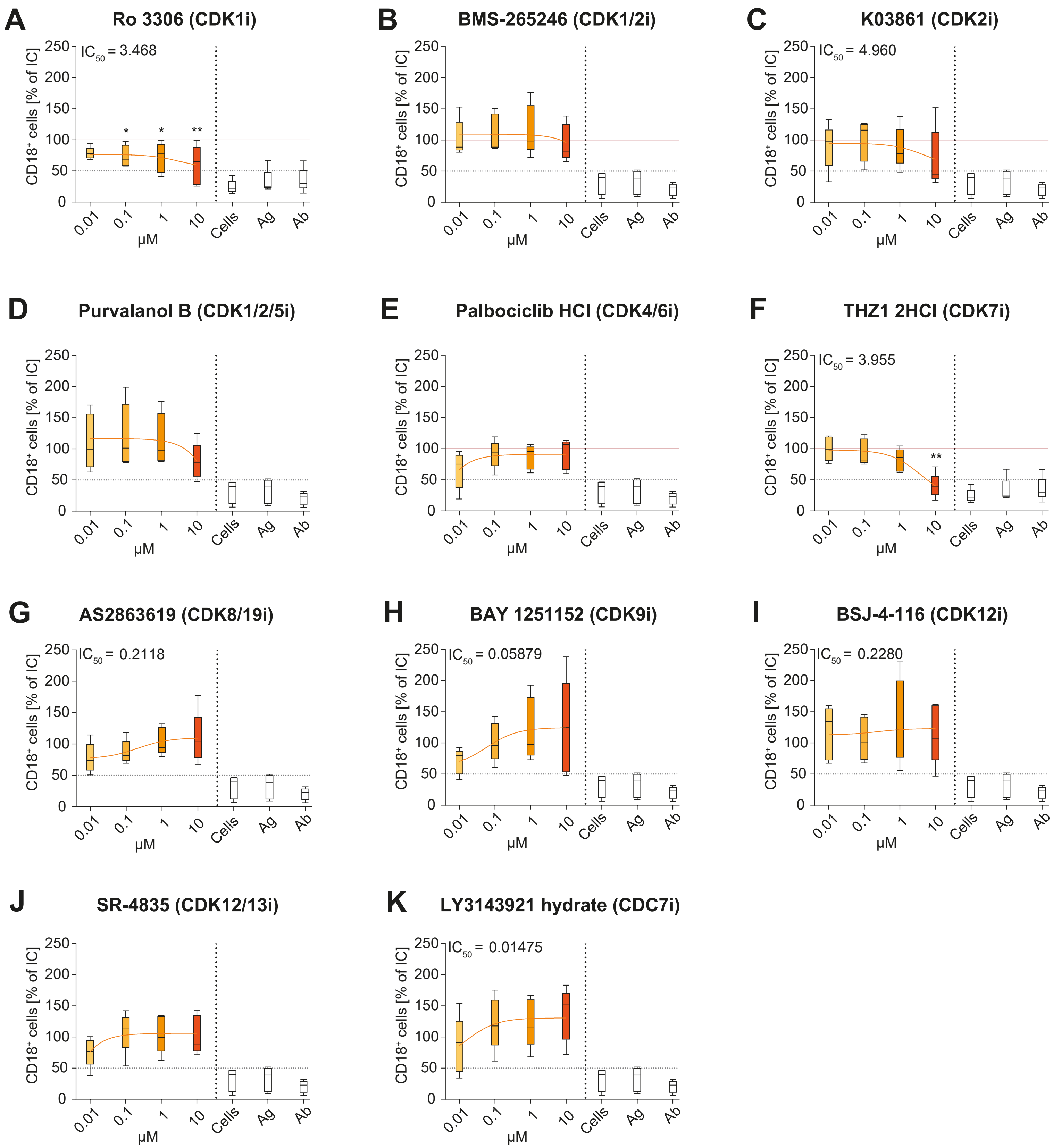

### Fig. S3

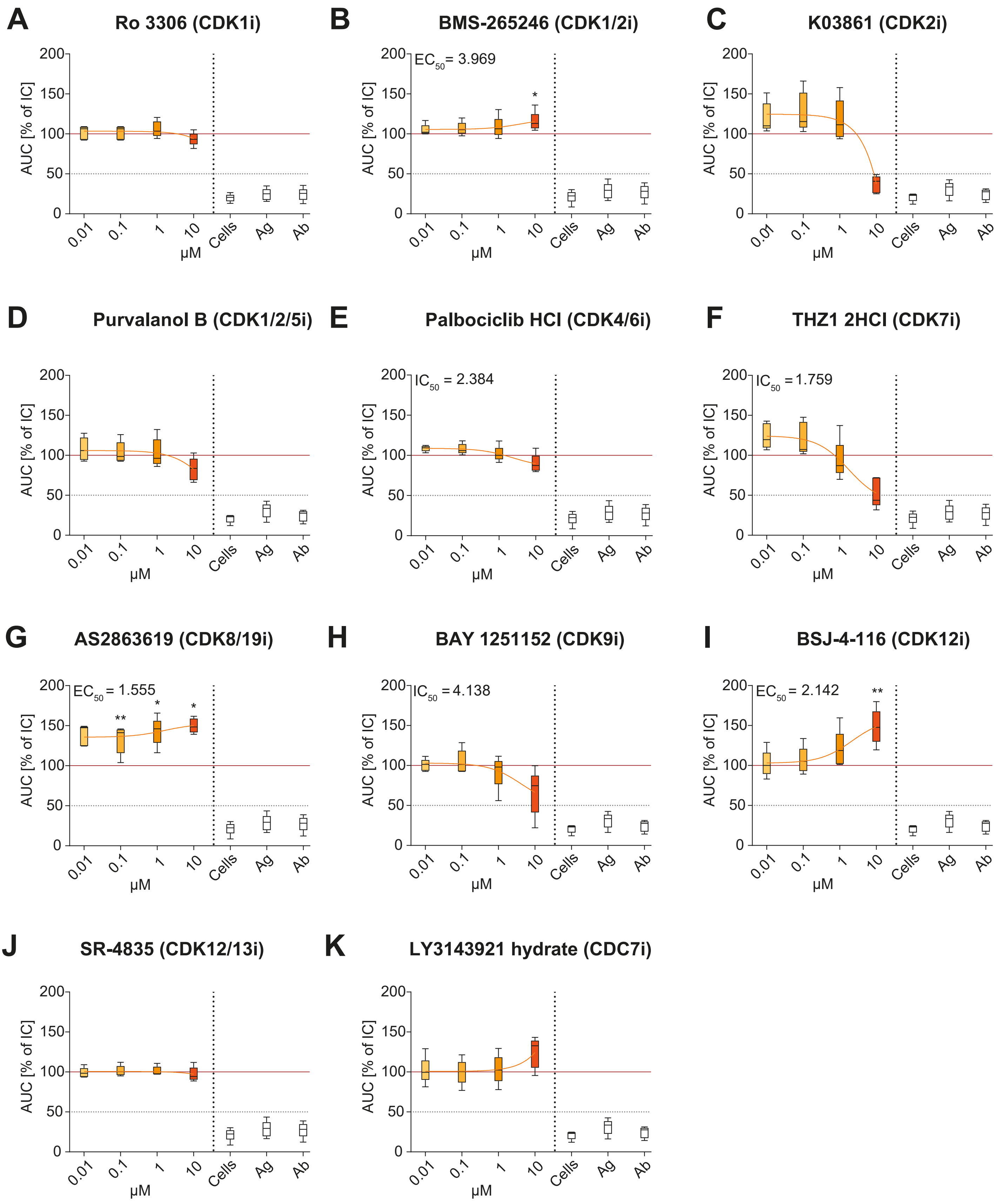

### Fig. S4

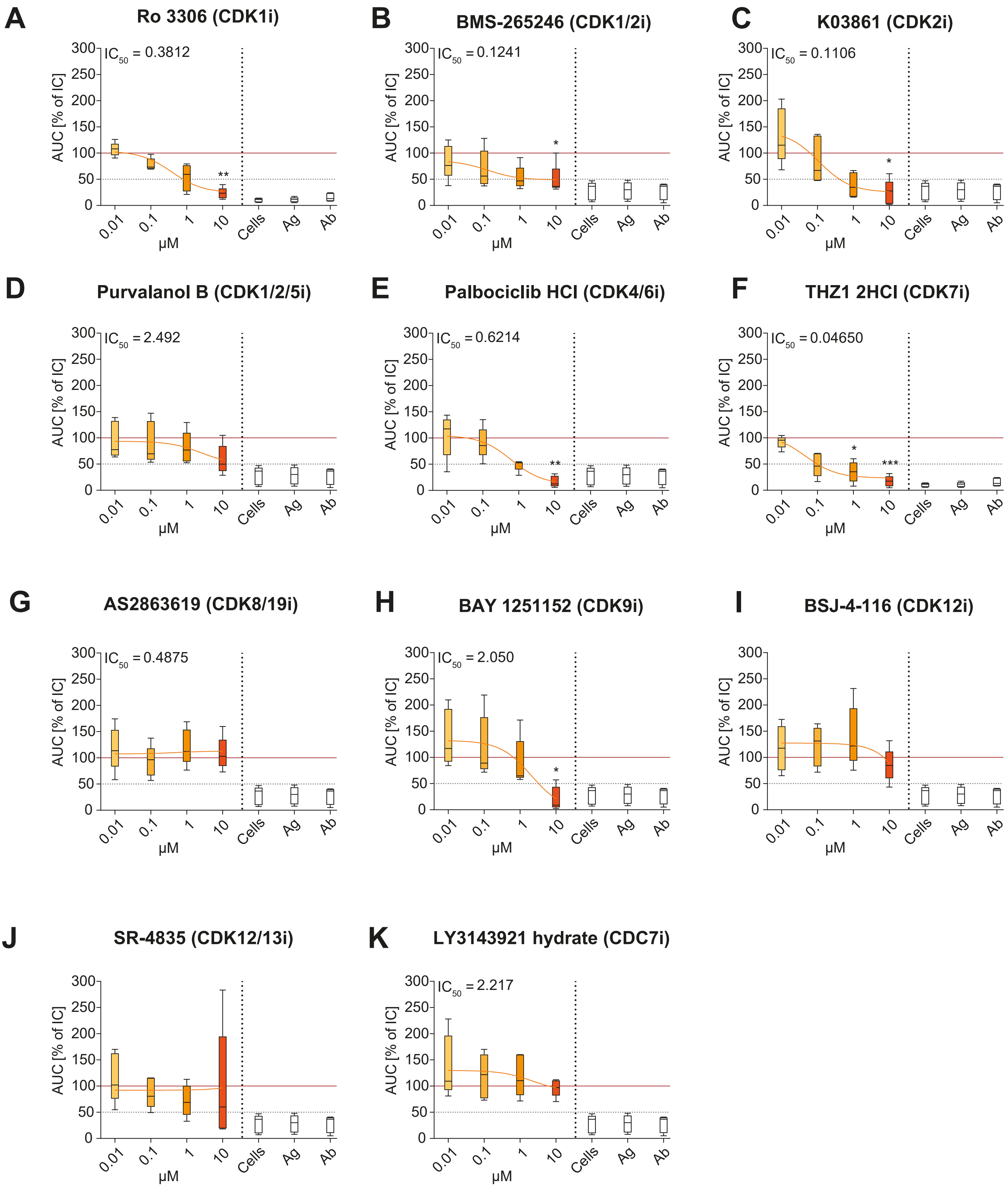

### Fig. S5

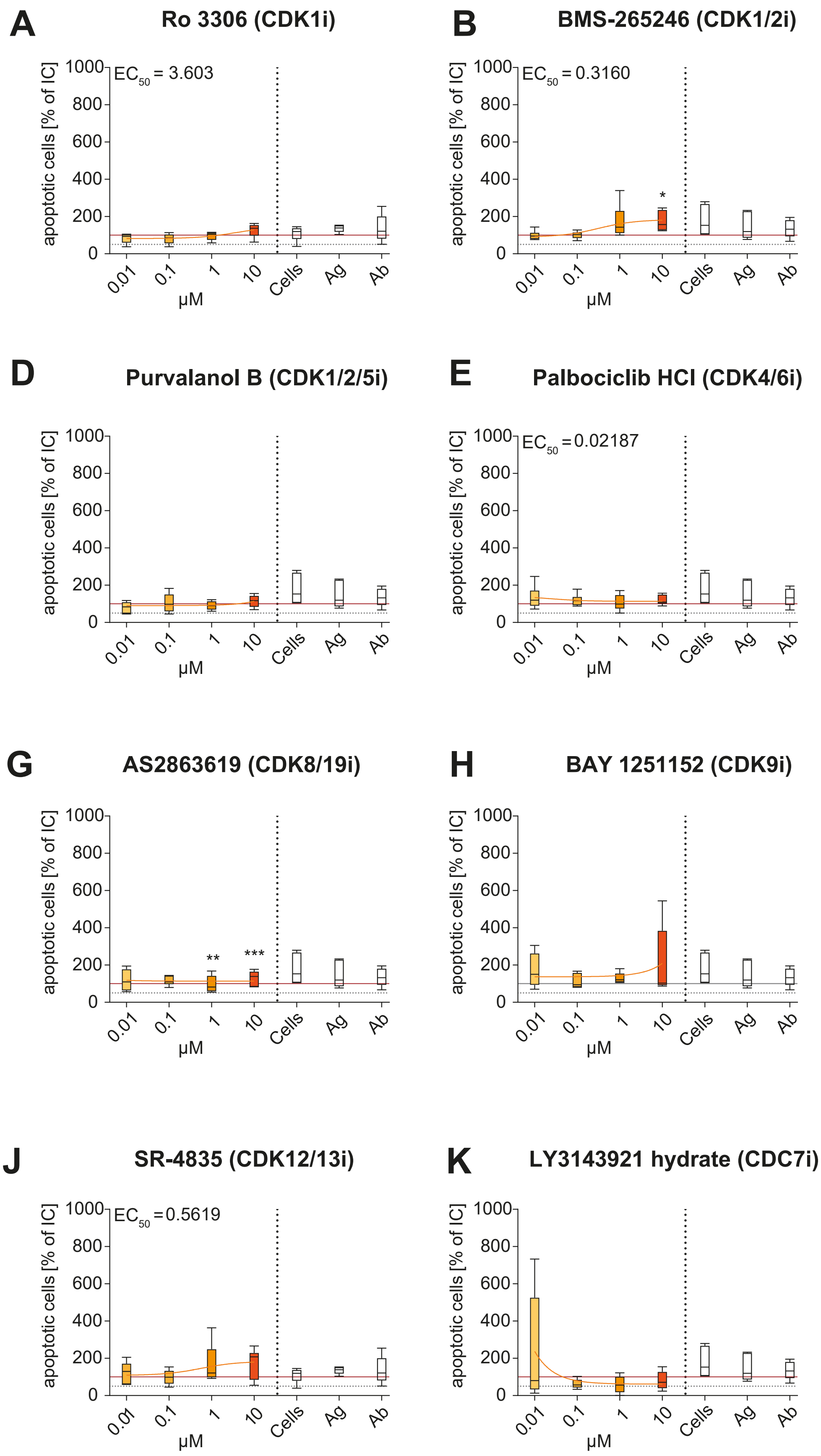

### Fig. S6

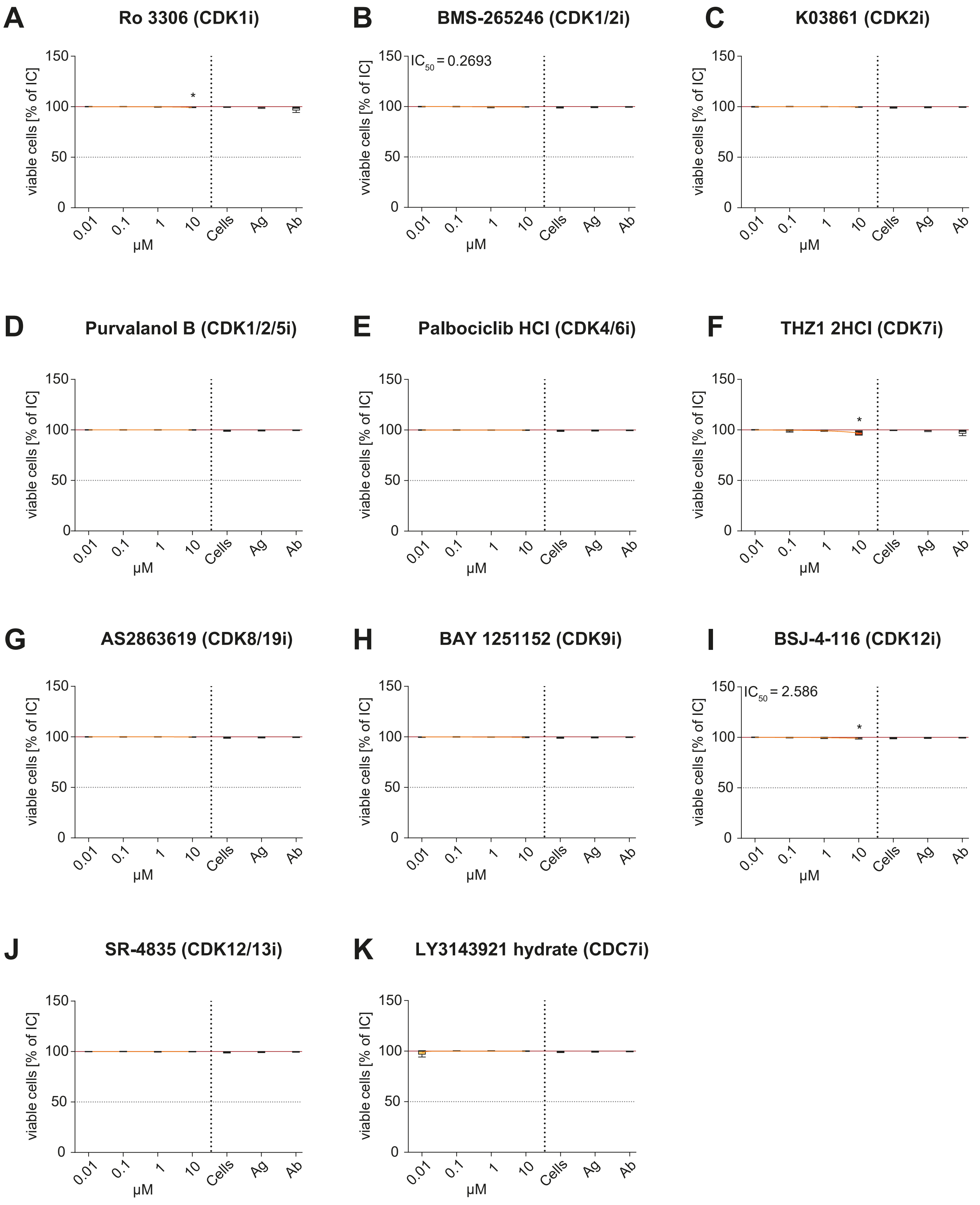

### Fig. S7

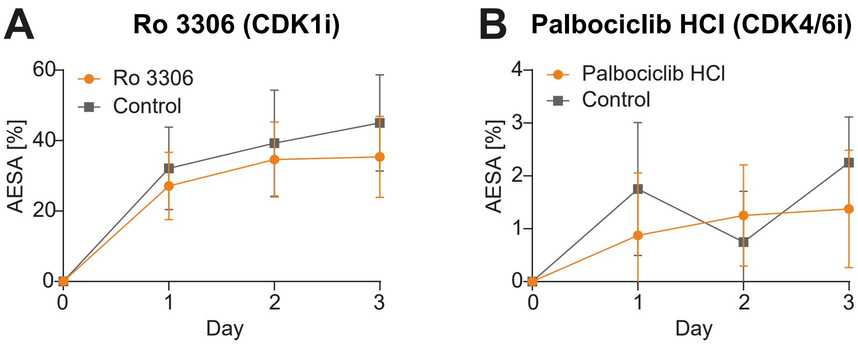
